# A DOMON domain-cytochrome b561 protein acts as a ferric reductase in iron homeostasis and impacts primary root growth under phosphate deficiency

**DOI:** 10.1101/2023.10.10.561639

**Authors:** Joaquín Clúa, Jonatan Montpetit, Yves Poirier

**Affiliations:** Department of Plant Molecular Biology, Biophore Building, University of Lausanne, 1015 Lausanne, Switzerland

## Abstract

Iron (Fe) and phosphate (Pi) are essential nutrients for plant growth. Several interactions between Fe and Pi homeostasis have been described, such as the Fe-dependent inhibition of primary root growth under Pi deficiency. This response involves the formation of apoplastic Fe^+3^-malate complexes in the root meristem which implicates the oxidation of Fe^+2^ by the LPR1 ferroxidase. However, how is the reduced Fe^+2^ generated in the root meristem and the Fe^+3^/Fe^+2^ ratio regulated is unknown. Here, we have identified a gene in *Arabidopsis thaliana*, named *CRR*, that is implicated in primary root growth under Pi deficiency. Under low-Pi conditions, the *crr* mutant showed an enhanced reduction of primary root growth that was associated with increased accumulation of apoplastic Fe in the root meristem and a reduction in meristematic cell division. Conversely, CRR overexpression rendered primary root growth insensitive to low-Pi inhibition, reduced root apoplastic Fe deposition, and impacted the expression of genes involved in Fe and redox homeostasis. CRR is a member of an uncharacterized CYBDOM protein family possessing a cytochrome b561 (CYB561) with an N-terminal DOMON domain. We demonstrated that CRR localizes to the plasma membrane and possesses ascorbate-dependent ferric reductase activity. The *crr* single mutant and the *crr hyp1* double mutant, which harbored a null allele in another member of the CYDOM family, showed increased tolerance to high-Fe stress upon germination and seedling growth. In contrast, CRR overexpression was associated with increased uptake and translocation of Fe to the shoot and resulted in plants highly sensitive to Fe excess toxicity. Our results thus identify a ferric reductase implicated in root Fe acquisition and homeostasis and reveal a biological role for CYBDOM proteins in plants.

## Introduction

Phosphorus (P) and iron (Fe) are both essential elements for plants. P is found in cells in the form of orthophosphate (PO_4_^−3^) and is included in nucleic acids, proteins, lipids, and numerous other small molecules playing essential roles in energy metabolism, such as adenosine triphosphate (ATP) (Poirier et al., 2022). Fe cycles between two oxidation states, Fe^+3^ (ferric) and Fe^+2^ (ferrous), and participates in numerous fundamental processes and pathways involving redox reactions, including respiration and photosynthesis (Connorton et al., 2017). Although typically abundant in most soils, both P and Fe are largely present in insoluble forms, with P often forming complexes with Fe (hydro)oxides. In most soils, the amount of soluble inorganic phosphate (Pi; H_2_PO_4_^−^ or HPO4^−2^), the form of P acquired by plant roots, is typically 0.5–10 µM, while the concentration of soluble Fe in well-aerated soils can be as low as 100 pM in calcareous soils (Marschner, 1995; Frossard et al., 2000). In waterlogged soils with an acidic pH and low redox potential, as are often encountered in many wetland rice (*Oryza sativa*)-growing areas, high levels of Fe^+2^ can lead to iron toxicity (Onaga et al., 2016). In all plants, except members of the Gramineae, the main strategy to acquire Fe through the roots involves three main steps: 1) Fe^+3^ solubilization from the soil via acidification, implicating H^+^-ATPase pumps such as AHA2, as well as the release of low molecular weight organic compounds capable of chelating Fe^+3^, such as malate, flavins, and coumarins; 2) the reduction of soluble Fe^+3^ to Fe^+2^ by the plasma membrane ferric reductase FRO2; and 3) the import of Fe^+2^ via the IRT1 transporter (Kobayashi and Nishizawa, 2012; Brumbarova et al., 2015; Tsai and Schmidt, 2017; Robe et al., 2021). Similarly, the acquisition of Pi from P–Fe insoluble complexes also involves Pi solubilization via medium acidification and the release of organic acids (citrate, malate), followed by H^+^–Pi co-transport by PHT1 transporters (Nussaume et al., 2011).

Numerous interactions have been described in the pathways involving Fe and Pi homeostasis (Hanikenne et al., 2021); for example, Pi deficiency was shown to result in an increased plant Fe content (Misson et al., 2005; Hirsch et al., 2006; Ward et al., 2008) and the downregulation of numerous genes involved in Fe acquisition, such as *IRT1* and *FRO2*, while at the same time increasing the expression of other genes, such as *FER1* and *YSL8*, implicated in Fe storage and transport, respectively (Misson et al., 2005; Müller et al., 2007; Thibaud et al., 2010; Lan et al., 2012). Conversely, Fe deficiency can result in Pi accumulation (Zheng et al., 2009; Saenchai et al., 2016). Strong evidence that this pattern of co-regulation is not simply the result of the increased availability of Fe under Pi-deficient conditions, and vice-versa, comes from the implication of two central regulators of Pi and Fe homeostasis. PHR1 and its close homolog PHL1 are members of the GARP family of transcription factors and are responsible for the majority of the transcriptional changes occurring under Pi deficiency (Bustos et al., 2010). Several of the Fe-homeostasis genes activated by Pi deficiency, such as *FER1*, are direct targets of PHR1 (Bournier et al., 2013; Briat et al., 2015). In the Fe pathway, members of the Brutus family of E3 ligases (BTS, BTSL1, and BTSL2 in *Arabidopsis thaliana* and HRZ, HRZ2, and HRZ3 in rice) participate in the degradation of key transcription factors involved in Fe homeostasis, such as FTI in *A. thaliana* and PRI in rice, and are also thought to act as Fe sensors (Kobayashi et al., 2013; Selote et al., 2015; Zhang et al., 2017; Rodriguez-Celma et al., 2019; Zhang et al., 2020). A recent study in rice has shown that HRZ1 and HRZ2 interact with and mediate the degradation of PHR2 (the rice orthologue of PHR1), explaining why Fe deficiency can attenuate the Pi deficiency response (Guo et al., 2022). Furthermore, the same study found that PHR2 transcriptionally downregulates the expression of the HRZs. PHR1/2 and BTS/HRZ thus form a reciprocal inhibitory module contributing to the coordination of Pi and Fe homeostasis.

Interactions between Pi and Fe also influence root growth (Bouain et al., 2019). While primary root growth is inhibited under either Pi or Fe deficiency, combined Fe and Pi deficiency results in a reversion of this growth suppression. Genetic screens involving mutants in the response of primary root growth under Pi deficiency have identified several genes involved in the Fe–Pi interaction during root growth. *LPR1* encodes a ferroxidase that converts Fe^+2^ to Fe^+3^, while *PDR2* encodes an endoplasmic reticulum (ER)–localized P5-type ATPase thought to negatively affect LPR activity via an unknown mechanism (Ticconi and Abel, 2004; Svistoonoff et al., 2007; Ticconi et al., 2009; Naumann et al., 2022). While the primary root growth of the *lpr1* mutant is unaffected by Pi deficiency, the *pdr2* mutant is hypersensitive under the same condition (Ticconi et al., 2004; Svistoonoff et al., 2007; Ticconi et al., 2009). Additional components of the pathway include the malate efflux channel ALMT1; the STOP1 transcription factor, which regulates *ALMT1* expression; the ALS3 and STAR1 subunits, which together form a tonoplast ABC transporter complex; and the CLE14 peptide receptors CLV2 and PEPR2 (Balzergue et al., 2017; Dong et al., 2017; Gutierrez-Alanis et al., 2017; Mora-Macias et al., 2017). The current model proposes that, under Pi-deficient conditions, malate secretion mediated by ALMT1 and STOP1 chelates Fe^+3^, allowing its accumulation in the apoplast of the root meristem and elongation zones, where it inhibits cell elongation and division. Apoplastic Fe^+3^ would participate in a redox cycle implicating the LPR1 ferroxidase, generating reactive oxygen species (ROS) and leading to a reduction in cell-to-cell communication via callose deposition, meristem differentiation by the activation of the CLE14–CLV2/PERP pathway, and a reduction in cell wall extensibility by the action of peroxidases (Müller et al., 2015; Balzergue et al., 2017; Gutierrez-Alanis et al., 2018). Blue light has also been implicated in this pathway, both as a cryptochrome-mediated signal transmitted to the roots and as a catalyst in Fenton reactions (Zheng et al., 2019; Gao et al., 2021). Despite these insights, the nature of the protein(s) mediating the Fe^+3^ to Fe^+2^ reduction involved in the proposed redox cycle is yet to be elucidated (Abel, 2017; Gutierrez-Alanis et al., 2018; Crombez et al., 2019).

We recently identified the ER chaperones Calnexin1 (CNX1) and Calnexin2 (CNX2) as novel components of the Fe-dependent inhibition of primary root growth under Pi deficiency (Montpetit et al., 2022). A proteomic analysis of the *cnx1 cnx2* double mutant identified a reduction in the amount of an uncharacterized CYB561 protein with a N-terminal DOMON domain, hereafter named CRR. Here, we show that CRR encodes a novel plasma membrane– localized ferric reductase that participates in Fe homeostasis as well as in the control of primary root growth under Pi deficiency.

## Results

### *CRR* encodes a CYBDOM protein that is less abundant in the *cnx1 cnx2* double mutant

In a previous work, we showed that the *A. thaliana cnx1 cnx2* double mutant showed a reduction in primary root growth under Pi deficiency, which was associated with an increased apoplastic Fe accumulation at the root meristem (Montpetit et al., 2022). Since the CNX proteins are components of the ER protein folding and quality control machinery, we hypothesized that the root phenotype of the *cnx1 cnx2* mutant could be a consequence of the misfolding, degradation, and/or destabilization of a particular set of proteins. To test this hypothesis, we performed a proteomic analysis of Col-0 and *cnx1 cnx2* mutant roots grown for 7 days in Pi-sufficient or -deficient conditions (+Pi or –Pi, respectively). In the +Pi condition, there were only 18 differentially abundant proteins between Col-0 and the *cnx1 cnx2* mutant, whereas in the –Pi condition, there were 136 differentially abundant proteins (Supplementary Table 1 and Figure S1A). The higher number of differentially abundant proteins in the –Pi treatment likely reflects the fact that the *cnx1 cnx2* mutant only showed a root growth phenotype under Pi deficiency (Montpetit et al., 2022). A Gene Ontology (GO) analysis of the differentially abundant proteins revealed they were enriched in functions related to ER stress, protein repair, and the response to unfolded proteins, indicating a predominant effect on ER protein homeostasis in the *cnx1 cnx2* mutant (Figure S1B).

To identify candidate genes implicated in the primary root growth phenotype of the *cnx1 cnx2* mutant, we focused on proteins that were differentially abundant in both the +Pi and –Pi conditions. Among the seven candidates was a member of the CYBDOM family (AT3G25290) that we named CRR for <u>C</u>YBDOM <u>R</u>OOT <u>R</u>EDUCTION (Figure S1C). As a typical CYBDOM protein, CRR contains a signal peptide at its N-terminal end, followed by a DOMON domain (<u>IPR005018</u>) and a cytochrome b561 (CYB561) (<u>IPR028836</u>) (Figure 1A and S2A). CYBDOM proteins have unknown functions in plants, and the *A. thaliana* genome encodes 10 of them (Preger et al., 2009). Notably, the mouse and human genomes each contain a single CYBDOM protein, named SDR2, and expression of the mouse SDR2 in *Xenopus* oocytes revealed Fe^+3^ reductase activity (Vargas et al., 2003). As expected for CYBDOM members, the CYB561 domain of CRR contains five transmembrane alpha helices as well as four conserved histidine groups involved in coordinating two heme *b* molecules (Asard et al., 2013) (Figure 1A, S2A– B). The N-terminal DOMON domain is predicted to be located in the apoplast and contains a conserved methionine and histidine previously shown to be involved in coordinating another *b* heme group in the cellobiose dehydrogenase of the fungus *Phanerochaete chrysosporium* (Hallberg et al., 2000; Asard et al., 2013) (Figure 1A, S2A–B). CRR has previously been detected in proteomic experiments analyzing purified plasma membranes, as reported on the SUBA database (https://suba.live/). To validate these predictions, we first analyzed the subcellular localization of a CRR-GFP translational fusion protein expressed under the CaMV *35S* promoter in *A. thaliana*. The confocal microscopy images reveal that CRR is mainly localized at the plasma membrane with a small fraction at the ER (Figure S3A). To further validate this localization, we transiently expressed CRR-GFP in *Nicotiana benthamiana* leaves together with a plasma membrane (CBL1-OFP) (Batistič et al., 2010) or ER (ER-rk-RFP) (Nelson et al., 2007) marker. CRR-GFP mostly co-localized with the plasma membrane marker, including during plasmolysis, with a small fraction associated with the ER marker, suggesting that the protein transits the secretory pathway from the ER to the plasma membrane (Figure S3B).

**Figure 1.**
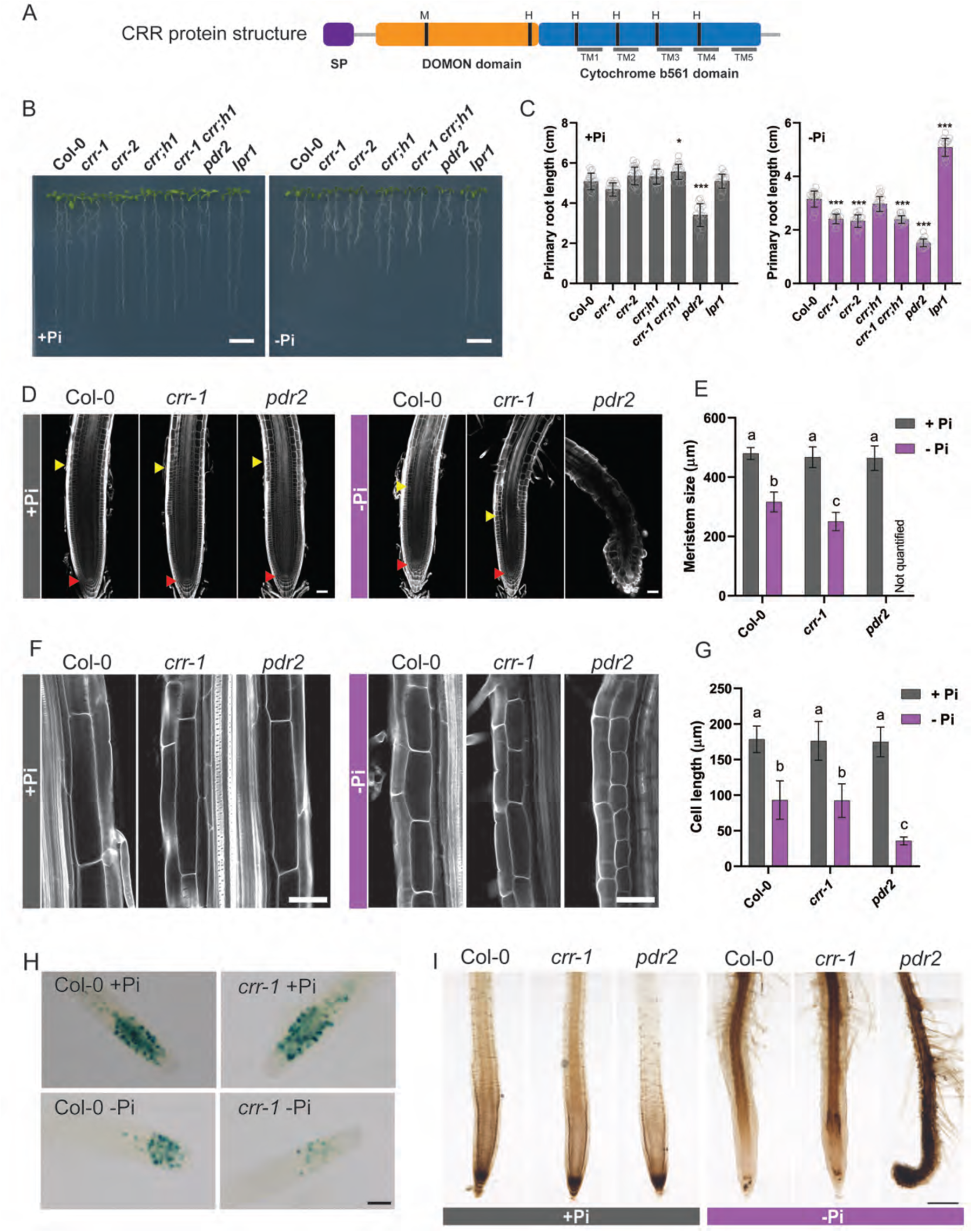
*CRR* encodes a CYBDOM protein involved in primary root growth under phosphate deficiency. **A**, Diagram of the CRR protein structure showing the positions of the signal peptide (SP), the DOMON and cytochrome b561 domains, the five transmembrane (TM) alpha helices, and the highly conserved methionine (M) and histidine (H) residues. **B**, Phenotypes of Col-0, *crr-1*, *crr-2*, *crr;h1*, *crr-1 crr;h1*, *pdr2,* and *lpr1* seedlings grown for seven days in Pi-sufficient (+Pi) or Pi-deficient (–Pi) conditions. Scale = 1 cm. **C**, Quantification of the primary root lengths of Col-0 and the different mutants at seven days after germination in +Pi and –Pi. The data are presented as means ± sd (n ≥ 15). The statistical differences were analyzed using a one-way ANOVA followed by Tukey’s multiple comparisons test. Significant differences to Col-0 are indicated (\*\*\**P* < 0.001; \*\**P* < 0.01; \**P* < 0.05). **D**, Root apical meristem (RAM) structure of Col-0, *crr-1,* and *pdr2*. Red arrows indicate the quiescent center and yellow arrows indicate the end of the meristematic zone. **E**, Quantification of the size of the RAM. **F**, Representative images showing the lengths of the cells in the root differentiation zone. In (D) and (F), the root cell walls were stained with Calcofluor White and visualized using confocal microscopy. Scale = 50 µm. **G**, Quantification of the lengths of the differentiated root cells of plants grown in +Pi and –Pi. In (E) and (G), the data are presented as means ± sd (n ≥ 4). The statistical differences were analyzed using a two-way ANOVA followed by Tukey’s multiple comparisons test. Different letters indicate significant differences (*P* < 0.05). **H**, GUS staining of Col-0 and *crr-1* carrying the G2/M transition marker pCYCB1::GUS. Scale = 100 µm. **I**, Visualization of the iron distribution at the root tip, obtained using Perls-DAB staining of plants grown in +Pi and –Pi. Scale = 200 µm.

The topology of CRR was assessed using a split-GFP assay (Xie et al., 2017). The C terminus of CRR was fused to the 11^th^ beta-strand of the GFP protein (CRR-GFP11) and co-expressed with a cytosolic or secreted version of the remaining 10 GFP beta-strands (GFP1–10). GFP fluorescence was only observed when CRR-GFP11 was co-expressed with the cytosolic GFP1– 10, confirming that the CRR C terminus faces the cytosol (Figure S3C). Considering the five transmembrane domains of the CYB561 domain, the DOMON domain is thus apoplastic.

### CRR is a negative regulator of the primary root growth response to Pi deficiency

The *cnx1 cnx2* double mutant showed a stronger reduction of primary root growth in response to Pi deficiency than did Col-0 (Montpetit et al., 2022); therefore, we assessed the root phenotype of the *crr* mutants under the same condition. The *crr-1* (*Salk_202530*) and *crr-2* (*Salkseq_039879.2*) mutants contain a T-DNA insertion in the first intron and second exon, respectively. A qRT-PCR analysis confirmed that both mutants had strongly reduced *CRR* mRNA levels in comparison with the wild-type (WT) plants grown in +Pi or –Pi conditions (Figure S4A, B). Furthermore, full-length *CRR* cDNA was undetectable by PCR in these mutants, indicating that both of them are knockout mutants (Figure S4C).

The primary roots of *crr-1* and *crr-2* were compared with Col-0 and the previously characterized *lpr1* and *pdr2* mutants after 7 days of growth on –Pi and +Pi media (Ticconi et al., 2004; Svistoonoff et al., 2007) (Figure 1B, C). As previously reported, the *pdr2* mutant showed pleiotropic growth defects in both +Pi and –Pi conditions, including a strong inhibition of primary root growth in the –Pi condition. On the other hand, the *lpr1* primary roots were completely insensitive to Pi deficiency, producing roots of equal length to those grown in the +Pi medium. The *crr-1* and *crr-2* mutant roots were approximately 25% shorter than those of Col-0 when grown in –Pi medium, but no difference was observed in the +Pi medium (Figure 1B, C). Similar to the *cnx1 cnx2* double mutant and *pdr2*, the short-root phenotype of the *crr* mutants under –Pi was dependent on the presence of Fe in the medium, as the primary root growth of the *crr-1* mutant in a –Pi –Fe medium was similar to that of Col-0 (Figure S5).

The *A. thaliana* CYBDOM protein family is divided into four different clades (Preger et al., 2009). CRR belongs to a clade of six members, and its closest homologous gene is AT4G12980, which was named CRR homolog 1 (*CRR;H1*) (Figure S2C–D). The high degree of identity between these two proteins could imply functional redundancy. To test this hypothesis, we compared the primary root growth of the *crr-1* and *crr;h1* single mutants, the *crr-1 crr;h1* double mutant, and Col-0 under both +Pi and –Pi conditions. No significant differences were observed between *crr;h1* and Col-0 or between the *crr-1 crr;h1* double mutant and the *crr-1* single mutant, leading to the conclusion that CRR;H1 is not involved in the primary root responses to Pi deficiency (Figure 1B, C).

Given that *crr-1* and *crr-2* are both T-DNA insertional mutants, we wanted to rule out the possibility that the observed root phenotype was a consequence of changes in the expression of *CRR*-neighboring genes. For this reason, we decided to edit the *CRR* locus using CRISPR/Cas9 technology. After generating transgenic lines expressing two different sets of guide RNAs, we obtained two independent homozygous lines in which *CRR* was edited. Line 25.6 has an in-frame deletion of 1120 nucleotides, generating an ORF potentially encoding a peptide of 93 aa, whereas line 22.6 has a deletion of 729 nucleotides, generating a hypothetical truncated version of CRR lacking the whole CYB561 domain (Figure S6A). When lines 25.6 and 22.6 were grown in +Pi and –Pi media, we observed a reduction in primary root growth relative to Col-0 under the –Pi condition only (Figure S6B, C). Together, these data show that *CRR* contributes to the primary root growth response under Pi deficiency.

### The short-root phenotype of *crr* is a consequence of defects in the root apical meristem maintenance associated with a hyperaccumulation of Fe

Primary root growth is essentially determined by cell division in the apical meristem and cell elongation in the root elongation zone. An inspection of the meristem morphology of Col-0 roots grown in –Pi showed the typical reduction in meristem size and the loss of identity of the quiescent center (QC) compared to roots grown in +Pi (Figure 1D, E). The general organization and morphology of the meristem of *crr-1* was similar to the Col-0 roots when grown under –Pi, including a loss of QC identity; however, the reduction in meristem size was exacerbated, resulting in a 20.9% smaller meristem than that of Col-0. In contrast, the *pdr2* mutant showed a very strong disorganization of the entire meristematic region, with large swollen cells.

Inspection of the root differentiation zone showed a 50% reduction in cell length in the Col-0 roots grown on –Pi compared with those on +Pi (Figure 1F, G), while in *pdr2*, this response was more pronounced, resulting in cell lengths reaching just 38% of those in Col-0 under –Pi. By contrast, *crr-1* showed no significant differences in cell length compared with those of Col-0 in the +Pi or –Pi conditions (Figure 1F, G). The CYCB1::GUS cell cycle reporter cassette was introgressed into the *crr-1* background to visualize meristematic cell division activity (Figure 1H). As previously reported, the meristematic cell division activity of Col-0 plants grown in –Pi was reduced when compared with the +Pi treatment (Ticconi et al., 2004). However, cell division activity under the –Pi condition was even more reduced in *crr-1* compared with Col-0, while *ccr-1* cell division activity was not affected under the +Pi condition. Taken together, these results reveal that CRR acts as a positive regulator of root apical meristem division and maintenance under Pi deficiency.

As the hypersensitive primary root growth inhibition of the *crr* mutant under the –Pi condition is dependent on Fe, we next explored the distribution of Fe in the root tips of *crr-1* and *pdr2* using Perls-DAB staining. In the +Pi condition, all the tested genotypes showed the strongest Fe accumulation at the root cap, consistent with previous reports (Müller et al., 2015; Mora-Macias et al., 2017) (Figure 1I). Under Pi deficiency, Col-0 displayed a reduced level of Fe at the root cap and an accumulation at the QC and the stem cell niche (SCN), as well as at the elongation and differentiation zones. Notably, *crr-1* roots followed the same Fe distribution pattern as Col-0, albeit with higher intensity. In agreement with its stronger root growth inhibition, *pdr2* showed a high accumulation of Fe in all root zones under Pi deficiency (Figure 1I). Taken together, our results show that *crr* accumulates more Fe than Col-0 under Pi deficiency, reducing cell division in the meristem. Importantly, while Fe accumulation in the elongation zone of the *crr* mutant was higher than in Col-0, it did not affect cell elongation.

A qRT-PCR analysis revealed an induction of *CRR* expression in the shoots under –Pi, while expression in roots was unaffected by the Pi status (Figure S7A). For unknown reasons, all experiments aimed at analyzing the endogenous expression pattern of *CRR* using either promoter fragments ranging from 200 to 5000 bp fused to GUS or gene fragments including the *CRR* promoter and *CRR* genomic coding regions or cDNA fused to GFP failed to reveal GUS or GFP expression, respectively. These observations indicate that the regulatory context for *CCR* expression might be more complex than typically observed for most *A. thaliana* genes and that important features, such as higher-order chromatin organization or epigenetic signatures, may play an important role in *CRR* promoter activity that cannot be reproduced in a transgenic setting (Kumar et al., 2021; Oudelaar and Higgs, 2021). Single-cell transcriptomic data from *A. thaliana* roots revealed the expression of *CRR* in the QC and SNC (initials), as well as in differentiating cortical, endodermal, and lateral root cap cells (Wendrich et al., 2020) (Figure S7C–D). The co-expression of *CRR* with *pUB25* and *pSPT* further indicated the expression of *CRR* in the root meristem (Figure S7B) (Wendrich et al., 2017). To identify the cells of the root tip in which the expression of *CRR* is required to complement the *crr* mutant phenotype, a *CRR-GFP* translational fusion construct was expressed under the control of promoters active in various regions of the root, including the SCN, endodermis, vasculature, cortical cells, and root cap (Figure S8A). Confocal microscopy images confirmed that the *LOVE1*, *SCR*, *WOX5*, *WOL*, *CO2*, *PEP*, *ATL75*, *MYB36*, and *LPR1* promoters allowed expression of CRR-GFP in their corresponding root domains (Figure S8B). Only transgenic plants expressing *CRR-GFP* under the control of the *LPR1* promoter (*pLPR1::CRR-GFP*), enabling strong expression in the SCN, endodermis, and pericycle, were able to complement the *crr-1* mutant (Figure S8C). This result indicates that *CRR* expression at the SCN is key to ameliorating root growth inhibition under low Pi and that *CRR* could play an antagonistic role to *LPR1*.

### Primary root growth is not inhibited by Pi deficiency in plants overexpressing *CRR*

To assess the effects of *CRR* overexpression on root growth, we analyzed transgenic lines expressing a *CRR-GFP* translational fusion construct (CRR OE) under the control of the CaMV *35S* promoter in both Col-0 and the *crr-1* mutant. Under +Pi conditions, the ectopic expression of *CRR-GFP* did not cause any obvious phenotype regarding shoot growth or root development in the Col-0 or *crr-1* backgrounds; however, under –Pi, *CRR* overexpression in both Col-0 and *crr-1* made primary root growth completely insensitive to Pi deficiency (Figure 2A and B). In our experimental conditions, both the CRR OE line and *lpr1* showed similar primary root growth that is unaffected by the Pi supply (Figure 2A and B).

**Figure 2.**
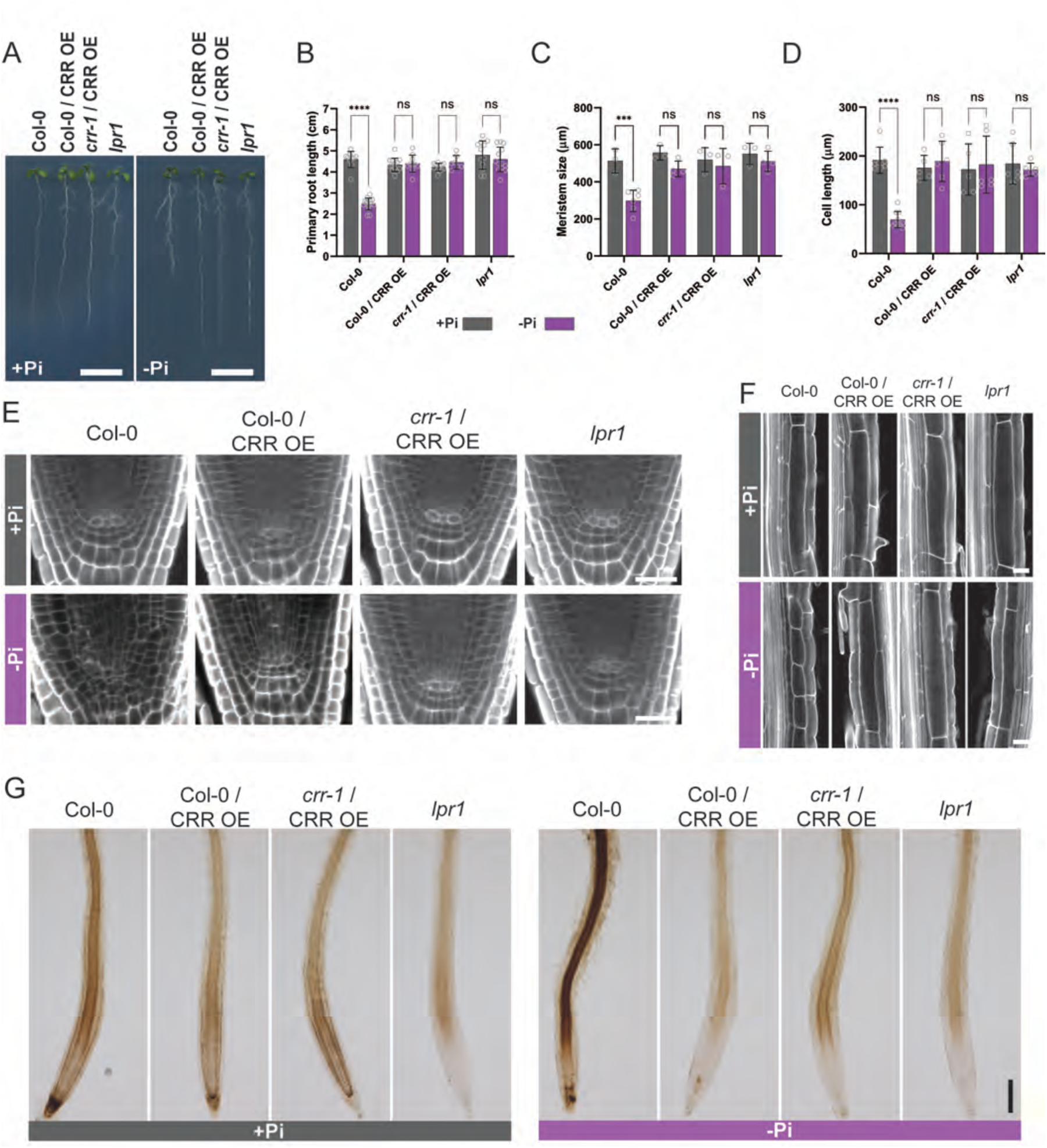
*CRR*-overexpressing plants are insensitive to phosphate deficiency–induced reductions in primary root growth. **A**, Phenotypes of the transgenic lines expressing *CRR-GFP* under the control of the *35S* promotor in the Col-0 (Col-0 / CRR OE) or *crr-1* (*crr-1* / CRR OE) backgrounds, as well as Col-0 and *lpr1,* grown on Pi-sufficient (+Pi) or Pi-deficient (–Pi) media for seven days. Scale = 1 mm. The roots of all lines were analyzed for **B**, primary root length; **C**, meristem size; and **D**, cell length in the differentiation zone. Data are presented as means ± sd (n ≥ 3) and, for each genotype, the statistical differences between treatments were analyzed using a two-way ANOVA followed by Šídák’s multiple comparisons test (\*\*\**P* < 0.001; \*\**P* < 0.01; \**P* < 0.05). **E**, Confocal microscopy images showing the quiescent center (QC) of the roots shown in (A). Scale = 20 µm. **F**, Pictures showing the lengths of the cells in the root differentiation zone. Scale = 25 µm. In (E) and (F), the root cell walls were stained with Calcofluor White and visualized using confocal microscopy. **G**, Visualization of the iron distribution at the root tip, obtained using Perls-DAB staining of plants grown in +Pi and –Pi. Scale = 200 µm.

Meristem size and the length of cells in the elongation zone of CRR OE were not significantly different in roots grown on +Pi or –Pi media, explaining its resemblance to the *lpr1* mutant (Figure 2C, D, F and S9). A closer inspection of the QC showed that, in Col-0 roots, the identity of the QC was lost under Pi deficiency and cells underwent differentiation, while the QC of the CRR OE roots grown in –Pi remained defined and undifferentiated as in the +Pi condition (Figure 2E). The same effect on the QC was observed for *lpr1*.

Analysis of Fe deposition by Perls-DAB staining showed that CRR OE had weaker Fe staining than Col-0 in the root cap under +Pi, but still more than *lpr1* (Figure 2G). In the –Pi condition, CRR OE and *lpr1* showed minimal Fe staining in the SCN and elongation zone. Altogether, these results suggest that CRR OE root growth is insensitive to Pi starvation as a consequence of a disruption in Fe accumulation at the root tip.

### *LPR1* and *PDR2* are epistatic to *CRR*

The genetic interactions between *CRR* and the main molecular players of the primary root responses to Pi deficiency were assessed by crossing *crr-1* with the long-root mutants *almt1* and *lpr1* and analyzing their root phenotypes under the +Pi and –Pi conditions. In +Pi, the root lengths of all lines were similar, with no obvious differences in terms of architecture (Figure 3A). In agreement with the macroscopic phenotype, no significant difference was found in the meristem size or cell length in the elongation zone between Col-0 and the mutant lines (Figure 3B). In –Pi, the macroscopic analysis of the root phenotypes revealed that the only fully epistatic interaction was between *lpr1* and *crr-1*, as the primary root length of *crr-1 lpr1* was not different from the *lpr1* single mutant (Figure 3A). In addition, *crr-1* showed partial epistasis with *altm1* for primary root length, with the *crr-1 altm1* double mutant producing a root length significantly lower than *altm1* but longer than *crr-1*. A microscopic analysis of the roots revealed that the meristem size and cell length in the elongation zone of the *lpr1* roots was not affected by the introgression of *crr-1*. In the case of *altm1*, the introgression of *crr-1* significantly reduced meristem division to a level similar to the *crr-1* single mutant but did not influence cell elongation (Figure 3B). These results further support the idea that CRR uncouples the meristem exhaustion process from cell elongation inhibition and explains why there is no complete epistasis between *crr* and *altm1*; *crr-1* is epistatic to *altm1* on its effect on meristem size under –Pi but has no effect on the length of cells in the elongation zone (Figure 3B).

**Figure 3.**
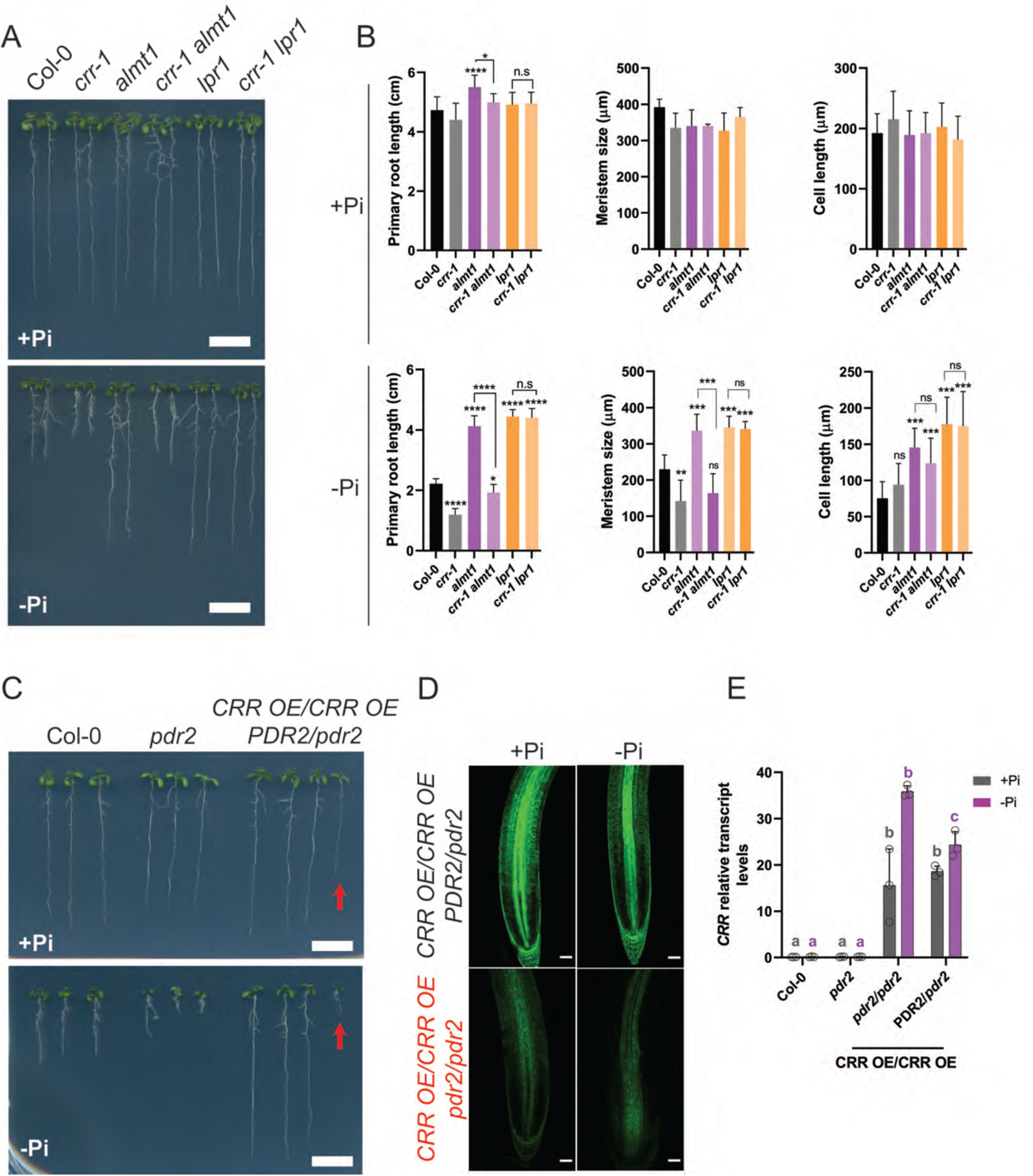
Genetic interactions between *CRR* and *ALMT1*, *LPR1*, *CLV2/PEPR2,* or *PDR2*. **A**, Primary root growth responses of plants grown in +Pi and –Pi media for seven days. Scale = 1 mm. **B**, Quantification of the primary root length, meristem size, and cell length in the differentiation zone of the roots of the different genotypes shown in (A). Data are presented as means ± sd (n ≥ 10 for root length, 3 for meristem size, and 10 for cell length). The statistical differences against Col-0 were analyzed using a one-way ANOVA followed by Dunnett’s multiple comparisons test. Additionally, selected comparisons and their statistical significance are indicated with brackets based on Tukey’s test. (\*\*\*\**P* < 0.0001; \*\*\**P* < 0.001; \*\**P* < 0.01; \**P* < 0.05). **C**, CRR OE and *pdr2-1* were crossed, and F_3_ seeds from a line homozygous for the CRR OE cassette but heterozygous for the *pdr2-1* allele were selected for analysis and compared with Col-0 and the parental *pdr2-1*. One quarter of the segregating population showed a short-root phenotype and were genotyped as homozygous *pdr2-1* (representative seedling highlighted by a red arrow), while all plants with a long-root phenotype were genotyped as either *PDR2*/*PDR2* or *PDR2*/*pdr2-1*. **D**, Confocal microscopy images showing *CRR-GFP* expression levels in the *PDR2*/*pdr2-1* or *pdr2-1/pdr2-1* backgrounds for plants grown in +Pi and –Pi media. Scale = 50 µm. **E**, Quantification of *CRR* transcript levels using qRT-PCR. The data represent the means ± sd (*n* = 3). The statistical differences between each genotype were analyzed using a one-way ANOVA followed by Tukey’s multiple comparisons test. Different letters indicate significant differences (*P* < 0.05).

To examine the genetic interaction between *CRR* and *PDR2*, the transgenic line CRR OE overexpressing *CRR-GFP* was crossed with the *pdr2-1* mutant. The F_3_ progeny from a line homozygous for the *CRR-GFP* transgene and heterozygous for *pdr2-1* were analyzed. The inspection of roots under –Pi showed that the F_3_ seedlings developed either a long-root phenotype resembling CRR OE or a hypersensitive short-root phenotype similar to *pdr2-1* in a ratio of 3:1 (Figure 3C). Genotyping showed that all seedlings with a short-root phenotype were *pdr2-1*/*pdr2-1* homozygous, while all seedlings with a long-root phenotype were *PDR2*/*PDR2* or *PDR*2/*pdr2-1*, thus demonstrating that *PDR2* is epistatic to *CRR*. PDR2 is localized in the ER, and the *pdr2-1* mutant is unable to maintain the integrity of the ER marker HDEL-GFP (Tecconi et al., 2009); therefore, we examined *CRR-GFP* expression levels in the *pdr2-1* mutant background. The results showed that the CRR-GFP protein levels were strongly reduced when introgressed in *pdr2-1*, while the mRNA levels remained unaffected, suggesting that a functional PDR2 is required for CRR protein stability (Figure 3D, E).

### The *crr* short-root phenotype under –Pi is enhanced by factors contributing to Fenton reactions

Fe-mediated Fenton reactions generating ROS have been implicated in the inhibition of primary root elongation under Pi deficiency, and such reactions are enhanced by low pH and H_2_O_2_ levels (Zheng et al., 2019). Primary root growth of Col-0, *crr-1*, and CRR OE grown in +Pi and –Pi media with different pH (5.2 to 6.0) (Figure S10A) or H_2_O_2_ contents (Figure S10B) was thus assessed. In +Pi, medium pH was found to result in similar root phenotypes in all the genotypes tested. In –Pi, however, the short-root phenotype of *crr-1* was greatly enhanced at low pH, while at higher pH, the *crr-1* root length was not different from Col-0 (Figure S10A). For all pH levels tested under –Pi, the primary root of the CRR OE line was longer than that of Col-0. Similarly, the addition of H_2_O_2_ in the +Pi medium had little influence on primary root growth, while it enhanced the reduction of *crr-1* root growth under –Pi (Figure S10B). These results support the hypothesis that the short-root phenotype of *crr-1* observed under –Pi depends on the amount of Fe accumulated at the SCN that can undergo Fenton reactions.

### CRR has ascorbate-dependent ferric reductase activity affecting root redox homeostasis

The potential implication of CRR acting in Fe^+3^ reduction was first tested by measuring ferric reductase activity in the roots of the CRR OE line using Fe-EDTA or FeCN as substrate. As a positive control, a line overexpressing the transcription factor bHLH39 (39OE) resulting in increased *FRO2* expression and high ferric reductase activity was used (Naranjo-Arcos et al., 2017). Our analysis showed that roots overexpressing *CRR* had a higher Fe-EDTA and FeCN reductase activity than Col-0 (Figure 4A). When compared with 39OE, the CRR OE roots showed a stronger reductase activity when using FeCN as an electron donor, but a significantly weaker reductase activity than 39OE when using Fe-EDTA. These findings suggest that CRR and the canonical FRO2 ferric reductase involved in Fe homeostasis have different affinities for various Fe–chelator complexes and might be involved in different processes. In agreement with this hypothesis, the primary root lengths of 39OE, *irt1*, and *fro2* grown in –Pi were similar to that of Col-0, indicating that neither FRO2 nor its associated Fe^+2^ transporter IRT1 influence the primary root growth response in – Pi conditions (Figure S11). Previous analyses have shown *FRO2* expression in the root hairs and epidermal cells of the mature root tissues but not in the root meristematic region (Connolly et al., 2003; Wu et al., 2005). Analysis of single-cell transcriptomic data of root tips shows a lack of *FRO2* expression in cells forming the root meristem (e.g., QC and SCN) (Figure S12).

**Figure 4.**
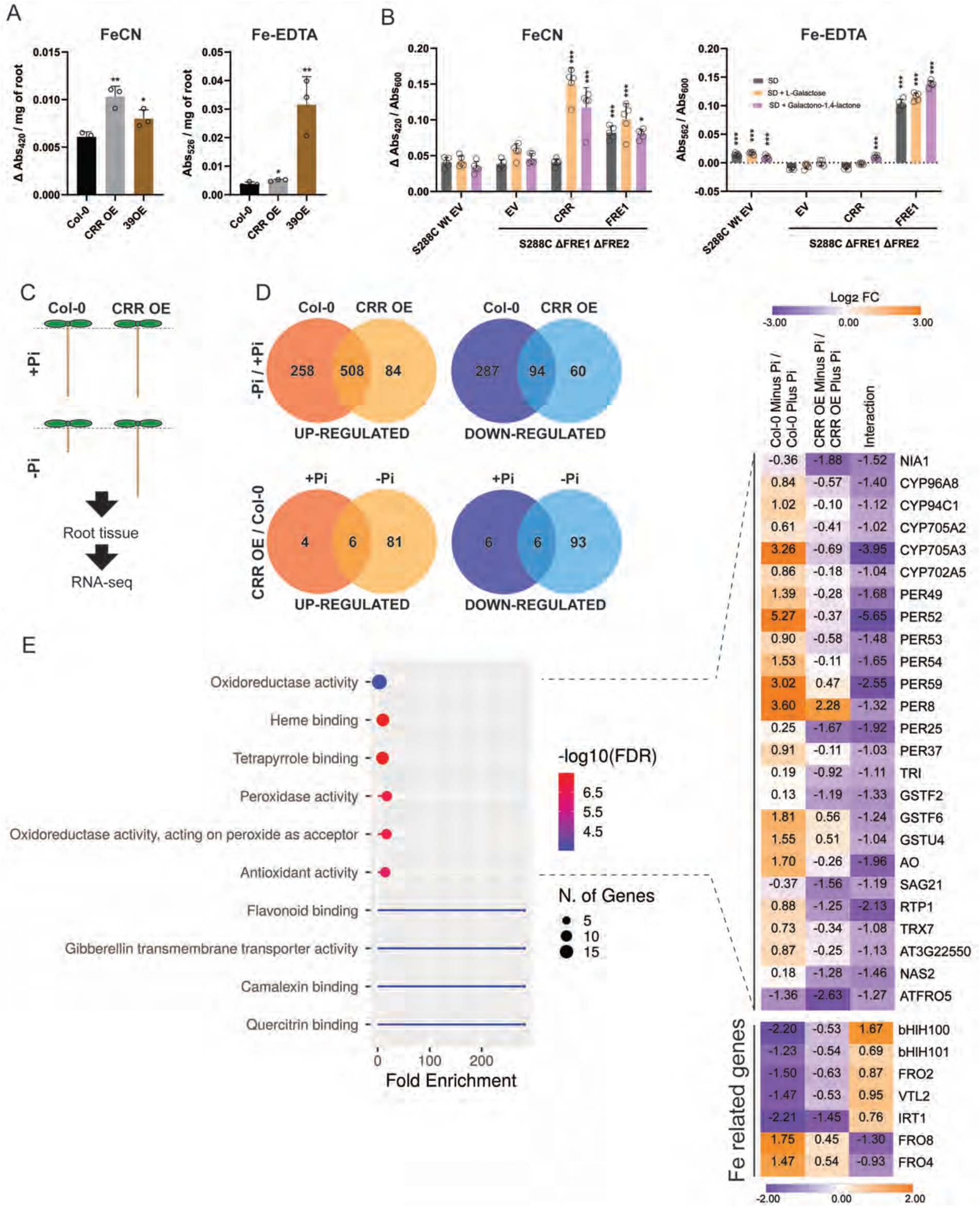
CRR has ferric reductase activity and its overexpression impacts the root transcriptome. **A**, Roots of Col-0, CRR OE, and 39OE (*bHLH39*-overexpressing line) were analyzed for their ferric reductase activity using either FeCN or Fe-EDTA as electron acceptors. FeCN iron reduction was measured as changes in absorbance at 420 nm and Fe-EDTA reduction was measured by the formation of ferrozine-Fe^+2^ complexes which absorb at 526 nm. The statistical differences were analyzed using a one-way ANOVA followed by Tukey’s multiple comparisons test. **B**, The *Saccharomyces cerevisiae* strain S288C or a mutant lacking the two main yeast ferric reductases FRE1 and FRE2 (*Δfre1 Δfre2*) were transformed with a plasmid driving the expression of *CRR* or *FRE1* under the control of the strong constitutive promotor *GPD*. The wild-type (WT) strain was transformed with an empty vector as a control (EV). All the strains were grown in synthetic defined media (SD) or SD supplemented with the ascorbate biosynthetic precursors L-galactose or galactono1,4-lactone. FeCN or Fe-EDTA were used as electron acceptors and iron reduction was quantified as in (A). The statistical differences from the EV control were analyzed using a two-way ANOVA followed by Tukey’s multiple comparisons test. In (A) and (B), data are presented as means ± sd (*n* ≥ 3; \*\*\**P* < 0.001; \*\**P* < 0.01; \**P* < 0.05). **C**, Schematic representation of the RNA-seq experimental design. Col-0 and CRR OE plants were grown for seven days in +Pi or –Pi media and root RNA was subjected to high-throughput sequencing. **D**, Venn diagrams showing the up-or downregulated differentially expressed genes (DEGs) of the treatment (–Pi / +Pi) or genotype (CRR OE / Col-0) responses. **E**, Gene ontology (GO) enrichment analysis of the downregulated DEGs found in the interaction between genotype and treatment. The lollipop graph shows the top 10 significantly enriched ‘molecular function’ GO terms. For each category, the fold enrichment, the number of genes, and the false discovery rate (FDR) are shown. The expression patterns of DEGs associated with redox process functions are displayed as a heatmap (top). The expression patterns of differentially expressed iron homeostasis–related genes are shown as a heatmap (bottom).

The Fe reductase activity of CRR was also assessed via its expression in the mutant *Saccharomyces cerevisiae* strain *Δfre1 Δfre2* lacking the two main endogenous ferric reductases, FRE1 and FRE2. As shown in Figure 4B, *CRR* expression did not enhance ferric reductase activity in the WT or *Δfre1 Δfre2* strains using either Fe-EDTA or FeCN. Some CYB561 proteins in animals and plants use ascorbate as an electron donor (Asard et al., 2013); however, *S. cerevisiae* is unable to biosynthesize ascorbate unless a chemical precursor like L-galactose or L-galacto-lactone is added to the medium (Spickett et al., 2000). When these precursors were added to the yeast growth medium, the expression of *CRR* in *Δfre1 Δfre2* resulted in a strong increase in ferric reductase activity with FeCN and a weaker increase with Fe-EDTA compared with the empty vector control (Figure 4B). Taken together, these results show that CRR is an ascorbate-dependent ferric reductase sensitive to different Fe chelation forms.

To assess how the high ferric reductase activity of CRR OE would affect its roots at the molecular level, in particular under –Pi where it displays a strong phenotype, a transcriptomic analysis of Col-0 and CRR OE roots grown under +Pi and –Pi was performed (Figure 4C). A dendogram of the 12 sequenced samples showed that the three replicates of each condition clustered together, confirming the good quality of the sampling (Figure S13A and Supplementary Tables 2 and 3). A principal component analysis (PCA) showed that in the +Pi samples, the transcriptomes of the Col-0 and CRR OE roots clustered together, while in –Pi, they clearly diverged, suggesting that *CRR* overexpression only significantly perturbs root gene expression under low-Pi conditions (Figure S13B). In agreement with this, an analysis of the differentially expressed genes (DEGs) between CRR OE and Col-0 revealed that only 22 genes were differentially expressed (10 upregulated and 12 downregulated) in +Pi, whereas in –Pi, there were 186 DEGs (87 upregulated and 99 downregulated, respectively) (Figure 4D, Figure S14A, and Supplementary Table 4).

An analysis of the DEGs between the –Pi/+Pi treatments revealed a large overlap of 508 genes among the upregulated genes in Col-0 and CRR OE. A GO analysis of this group showed that the main enriched terms were related to Pi starvation, Pi homeostasis, and Pi metabolism, suggesting that even though the CRR OE primary root is insensitive to the inhibitory effect of Pi deficiency on growth, the Pi deficiency signaling pathway was activated (Figure S14B). A detailed inspection of the canonical Pi starvation–responsive genes (e.g., *IPS1*, *SPX1*, *MGD2*, *PHT1;2*, and *PAP17*) indicated that all were upregulated to similar levels between Col-0 and CRR OE under Pi-deficient conditions (Figure S14C). Pi quantification in the roots and shoots showed that Pi levels in CRR OE and Col-0 were not significantly different, with a similar decrease in roots grown under –Pi conditions (Figure S14D). Overall, these data indicate that the CRR OE phenotype is restricted to primary root development under low-Pi conditions and that the Pi acquisition and Pi deficiency responses are not affected.

We analyzed the DEGs in the interaction between genotypes and treatments ((CRR OE –Pi / Col-0 –Pi) / (CRR OE +Pi / Col-0 +Pi)). A total of 190 DEGs were identified, of which 98 were upregulated and 92 were downregulated (Supplementary Table 4). In the set of genes downregulated in Pi-deficient CRR OE, a GO analysis based on molecular function showed a strong enrichment in categories related to redox homeostasis (e.g., oxidoreductase activity, peroxidase activity, and antioxidant activity) and Fe binding (heme and tetrapyrrole binding) (Figure 4E). A detailed analysis of these genes showed that several cytochromes p450, peroxidases, and glutathione *S*-transferases were deregulated in CRR OE (Figure 4E, upper heatmap). This finding prompted us to inspect the expression of genes more directly involved in Fe homeostasis. The *bHLH100* gene, encoding a key regulator of the Fe deficiency responses, was downregulated in the –Pi Col-0 roots; however, this inhibition was attenuated in the CRR OE line. Conversely, the upregulation of *FRO*8 in the –Pi Col-0 roots was attenuated in –Pi CRR OE (Figure 4E, lower heatmap). In addition, other Fe-related genes, such as *bHLH101*, *FRO2*, *VTL2*, *IRT1*, and *FRO4*, were mildly dysregulated in CRR OE. A GO analysis based on cellular component terms revealed that most of the DEGs in the interaction between genotypes and treatments were localized at the cell wall and/or in the apoplastic space (Figure S14E). Taken together, these results strongly suggest that CRR activity affects the redox status of the root apoplastic space as a consequence of changes in the amount of Fe accumulated in the apoplast, explaining the root growth insensitivity of CRR OE to –Pi, as well as the deregulation of the Fe-homeostasis genes.

### *CRR* expression levels affect *A. thaliana* tolerance to Fe stress

The findings that CRR has ferric reductase activity, that its overexpression causes the deregulation of Fe-homeostasis and redox-related genes, and that its expression level determines the amount of Fe accumulated at the root tip under –Pi conditions led us to speculate that CRR (and possibly other members of the CYBDOM family) might have a broader biological function in Fe homeostasis. Because the *crr-1* and CRR OE phenotypes are associated with conditions in which Col-0 roots hyperaccumulate Fe, we investigated the response of these lines to high Fe levels. We also included in the analysis the double mutant line *crr-1 hyp1*, which carries a knockout mutant allele of another member of the *A. thaliana* CYBDOM family, named *HYPERSENSITIVE TO PHOSPHATE 1* (*At5G35735*). In the accompanying manuscript, Giehl and collaborators show that HYP1 is also a ferric reductase involved in the growth response of primary roots to Pi deficiency by modulating the amount of Fe accumulated in the root, raising the possibility that these two genes have a partially redundant function.

Col-0, *crr-1*, CRR OE, and *crr-1 hyp1* were challenged to high amounts of Fe supplied as Fe-EDTA for 7 days from germination. Remarkably, CRR OE was hypersensitive to Fe stress, showing a mild to strong reduction in the fresh weight of its seedlings in media containing 400 to 600 µM Fe-EDTA, respectively (Figure 5A–C). Conversely, the *crr-1* and *crr-1 hyp1* mutants were more tolerant of the high-Fe stress in terms of the biomass generated. Similar results were obtained when seeds were sown in –Pi media with high Fe, with the exception that the Fe toxicity symptoms appeared at lower concentrations (350 µM Fe-EDTA), most likely because the removal of Pi enhances Fe accessibility via a reduction in the formation of insoluble Pi–Fe complexes (Figure S15A, B). Similar contrasting growth phenotypes were observed for the CRR OE and *crr-1 hyp1* plants grown for 7 days in a control basal MS medium and transferred into a medium with an additional 600 µM Fe-EDTA for 4 days (Figure S15C, D).

**Figure 5.**
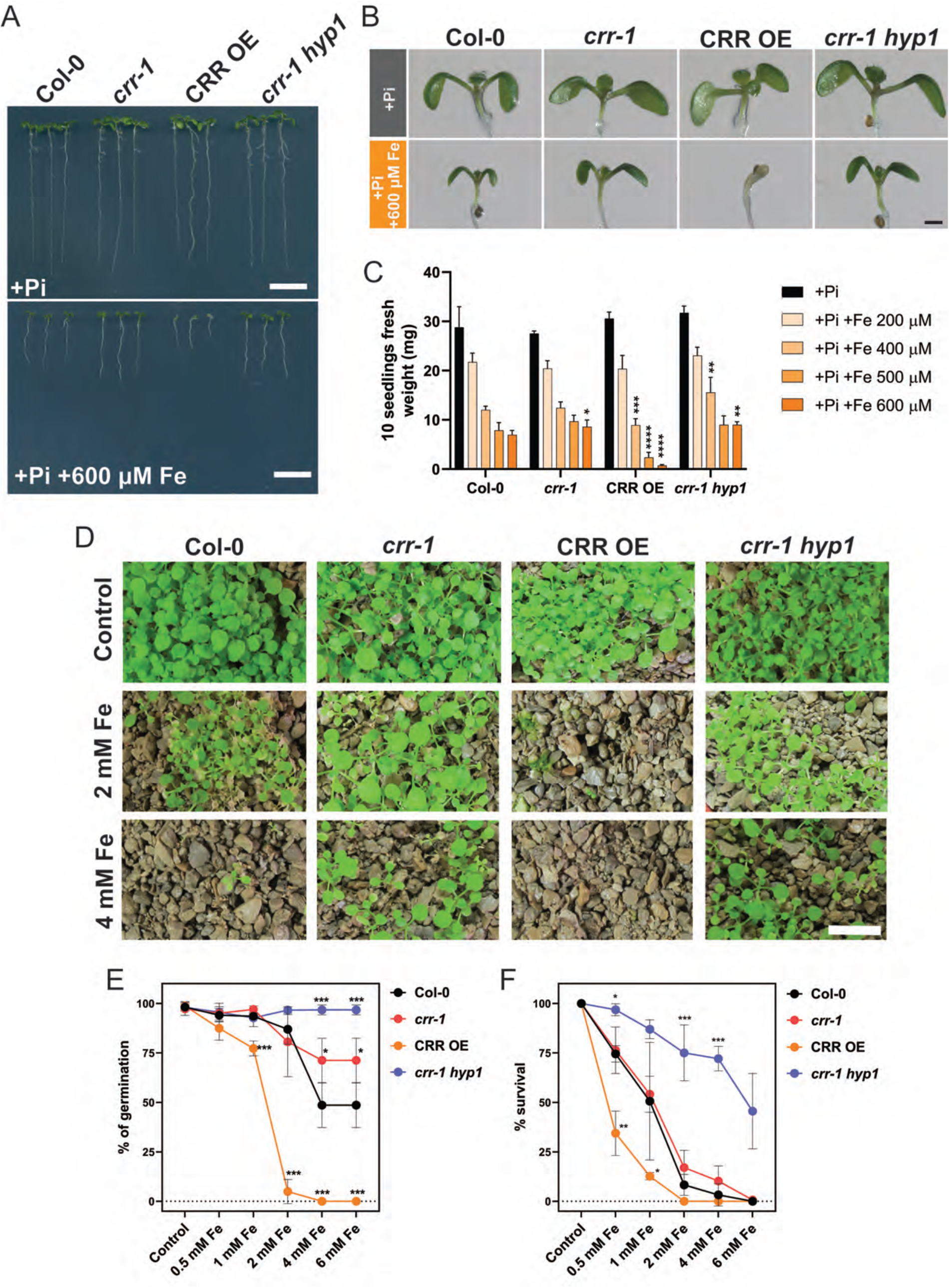
CRR is involved in iron toxicity tolerance. **A**, Phenotypes of Col-0, *crr-1*, CRR OE, and the double mutant *crr-1 hyp1* grown for seven days in the +Pi medium with or without an additional 600 µM Fe provided as Fe-EDTA. Scale = 1 mm. **B**, Magnification showing the shoot phenotype of the seedlings photographed in (A). Scale = 0.5 mm. **C**, Effect of increasing the concentration of Fe-EDTA on Arabidopsis seedling growth, determined by measuring the weight of 10 seedlings after 7 days of growth. **D**, Phenotypes of 15-day-old seedlings sown in a clay-based substrate watered with increasing concentrations of Fe-EDTA. Scale = 1 cm. **E**, Percentage of seeds that germinated in the clay-based substrate watered with increasing concentrations of Fe-EDTA, measured after five days. **F**, For the plants in (E) that had germinated by day 5, the percentage of plants that remained green and continued to develop for an additional 10 days was quantified. In (D), (E), and (F), the data are the means ± sd (*n* ≥ 3). The statistical differences within each condition were analyzed using a one-way ANOVA followed by Tukey’s multiple comparisons test. Significant differences from Col-0 are indicated (\*\*\**P* < 0.001; \*\**P* < 0.01; \**P* < 0.05).

The phenotypes associated with a high-Fe supply were also analyzed for the CRR OE and *crr-1*/*crr-1 hyp1* mutants grown under phototrophic growth conditions. In a clay-based substrate, Col-0 germination and growth were negatively affected when supplemented with a 2 mM Fe-EDTA solution, while CRR OE was unable to germinate under the same conditions (Figure 5D– E). Conversely, both the *crr-1* and *crr-1 hyp1* mutants were clearly more tolerant of Fe toxicity, showing higher levels of germination compared with Col-0 at Fe-EDTA concentrations of 4–6 mM (Figure 5D–E). Furthermore, of the seeds that germinated, the *crr-1 hyp1* mutant showed the highest level of survival after 15 days of stress, while CRR OE had the lowest survival rate (Figure 5F). The germination and survival rates of the *crr-1 hyp1* double mutant under high Fe were both higher than those of the single *crr-1* mutant (Figure 5E–F). Similar results were obtained when the experiments were performed with plants grown in peat-based soil (Figure S15E).

### CRR impacts Fe accumulation in the shoots

To understand the physiological mechanisms behind the high Fe–associated phenotypes of the *crr-1* / *crr-1 hyp1* mutants and CRR OE, the Fe distribution in the roots of seedlings grown in agar plates was examined. Perls-DAB staining of roots showed that *crr* and *crr hyp1* accumulated more Fe at the root tip than Col-0 under Fe stress conditions, whereas CRR OE accumulated less (Figure S16). Since the intensity of the staining reached near saturation, we performed a Perls staining without the DAB intensification step. This analysis revealed that while Col-0, *crr-1*, and *crr-1 hyp1* accumulated Fe at the QC and the root cap with only faint staining in the SCN under control conditions, CRR OE showed similar staining in the QC and SCN, but only very weak Fe accumulation in the root cap (Figure 6A). In the elongation and differentiation zones, Fe deposition was below the detection limits for all lines. In the medium with 350 µM Fe-EDTA, Col-0 showed Fe staining mainly in the QC and weakly in the SCN, a pattern similar to CRR OE under control conditions. In contrast to Col-0, *crr-1* and *crr-1 hyp1* displayed higher levels of Fe at the root meristem, including in the QC and SCN. There was an increase in Fe staining intensity in the elongation and differentiation zones of the *crr-1* single and *crr-1 hyp1* double mutants. In CRR OE, Fe was undetected in the root meristem and only a weak signal was obtained in the elongation and differentiation zones (Figure 6A).

**Figure 6.**
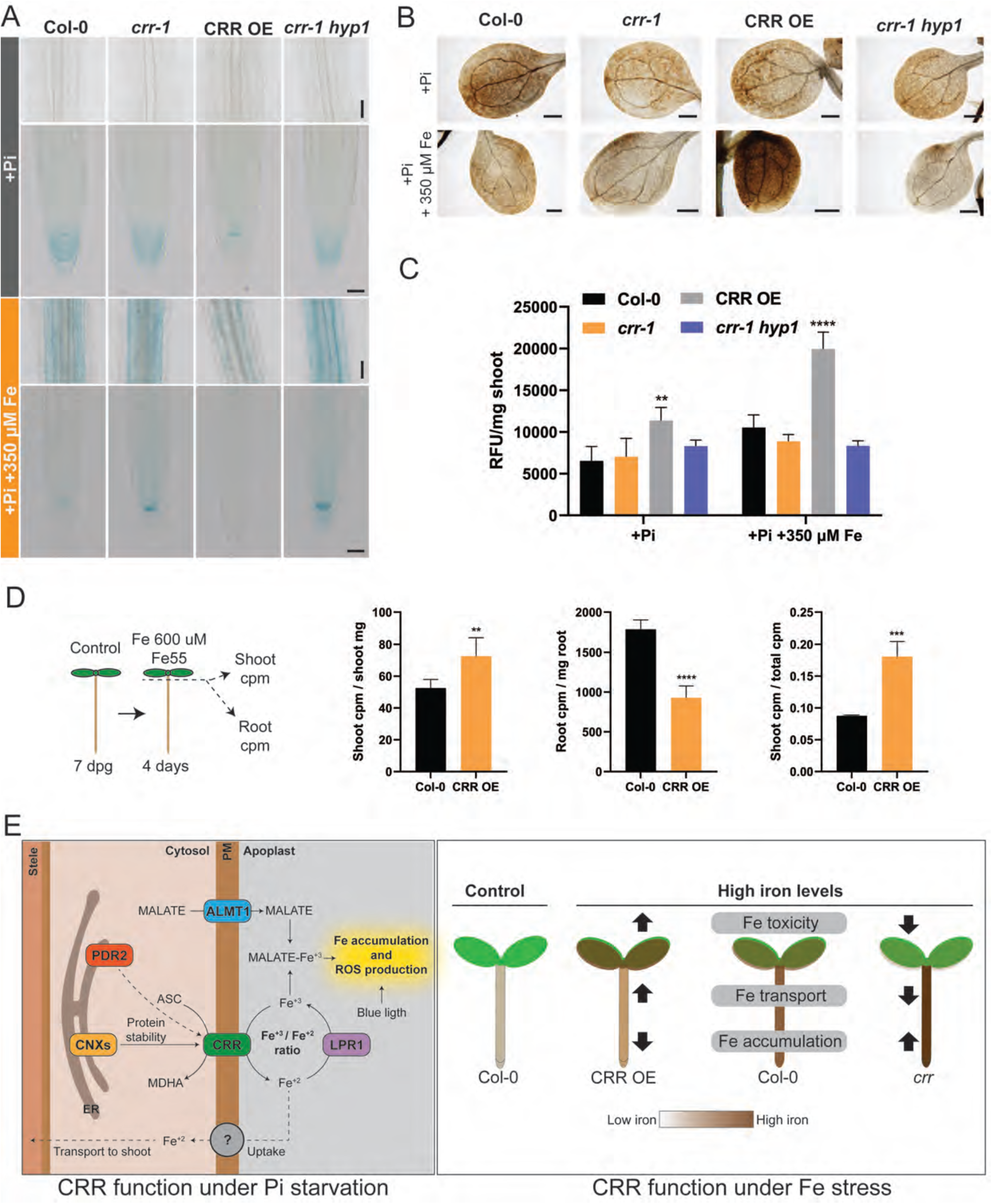
CRR impacts iron accumulation in the roots and its transport to the shoots. **A**, Phenotype of Col-0, *crr-1*, CRR OE, and *crr-1 hyp1* plants grown for seven days in the +Pi medium as a control, or in the +Pi medium supplemented with 350 µM Fe-EDTA. The roots were subjected to Perls staining to detect iron as a blue precipitate. Scale = 350 µM. **B**, Perls-DAB staining showing Fe accumulation patterns in the cotyledons. Plants were grown as in (A), and the shoots were subjected to Perls staining followed by a DAB intensification step. Scale = 500 µm. **C**, ROS in the shoots of plants grown as in (A), quantified using the fluorescence probe Carboxy-H2DCFDA and expressed as relative florescence units (RFU) per mg of tissue. The statistical differences within each condition were analyzed using a one-way ANOVA followed by Tukey’s multiple comparisons test. Significant differences from Col-0 are indicated (\*\*\**P* < 0.001; \*\**P* < 0.01; \**P* < 0.05). **D**, Schematic representation of the experimental design (left). Plants were grown for seven days in +Pi control conditions and transferred onto a +Pi medium containing 600 µM Fe-EDTA and 1.7 µCi per ml of ^55^Fe. After four days, the shoots and roots were collected separately, extensively washed, and the amount of ^55^Fe in the tissue was quantified as counts per million (cpm). Results are expressed as shoot cpm per mg of shoot, root cpm per mg of root, or the ratio shoot cpm over total plant cpm. The differences were statistically analyzed using an unpaired two-tailed t-test (\*\*\**P* < 0.001; \*\**P* < 0.01; \**P* < 0.05). In (C) and (D), the data are the means ± sd (*n* ≥ 4). **E**, Working model of CRR function under phosphate deficiency and iron stress. The model is explained in the Discussion.

Our analysis of Fe accumulation in the cotyledons using Perls-DAB staining showed that, when grown in a medium containing 350 µM Fe-EDTA, *crr-1 hyp1* accumulated less Fe than Col-0, whereas CRR OE accumulated more (Figure 6B). The possibility that Fe accumulation in the shoots of CRR OE was associated with increased ROS production was tested using carboxy-H2DCFDA. As shown in Figure 6C, the CRR OE shoots had higher levels of ROS when grown under both control conditions and with a 350 µM Fe-EDTA supply. Overall, an analysis of the Fe staining in both the roots and shoots revealed an antagonistic pattern between CRR OE and the *crr-1*/*crr-1 hyp1* mutants, with CRR OE accumulating more Fe in the shoots than the roots, resulting in ROS generation in the shoots. By contrast, the *crr-1* and *crr-1 hyp1* mutants accumulated more Fe in the roots and less in the shoots, suggesting that Fe reduction by CRR (and HYP1) may be associated with its transport from the roots to the shoots.

To further elucidate how CRR could modulate Fe transport, CRR OE seedlings grown for 7 days on a control medium were transferred into a medium containing 600 µm Fe-EDTA and supplemented with ^55^Fe for 4 days before measuring the amount of ^55^Fe in the roots and shoots. Consistent with the Perls-DAB staining, CRR OE contained a higher amount of ^55^Fe per milligram of tissue than Col-0 in the shoots and a lower amount in the roots, with a two-fold greater level of root-to-shoot ^55^Fe transfer than in the WT (Figure 6D). Altogether, these results suggest that *CRR* overexpression results in the greater accumulation of Fe in the shoots, affecting plant fitness under high Fe concentrations.

## Discussion

Studies on the response of primary roots to phosphate deficiency in *A. thaliana* have established a link between Fe deposition in the root tip apoplast with the reduction in meristematic cell division and cell elongation (Müller et al., 2015; Balzergue et al., 2017; Dong et al., 2017; Gutierrez-Alanis et al., 2017; Mora-Macias et al., 2017; Naumann et al., 2022). The central role of *LPR1*, encoding a ferroxidase, in apoplastic Fe accumulation and root growth under Pi deficiency implies the action of a ferric reductase to provide Fe^+2^ in the root meristem (Müller et al., 2015; Naumann et al., 2022). In *A. thaliana*, FRO2 is the primary root ferric reductase involved in Fe import into plants (Robinson et al., 1999). FRO2 is localized to the plasma membrane and primarily expressed in the root epidermis and root hairs (Connolly et al., 2003; Wu et al., 2005; Martin-Barranco et al., 2020). Although FRO2 paralogues are present in other subcellular membranes (e.g., FRO3 and FRO7 in chloroplasts and mitochondria, respectively), FRO2 remains the only FRO family member known to be involved in Fe reduction in the apoplast (Jain et al., 2014). While some coumarins secreted under Fe-deficient conditions at high external pH exhibit intrinsic Fe-reducing properties, their role in Fe acquisition at neutral and more acid pH levels appears negligible (Grillet and Schmidt, 2017; Rajniak et al., 2018; Tsai et al., 2018). Furthermore, even under alkaline conditions, Fe mobilized by coumarins requires the action of FRO2 for its acquisition (Fourcroy et al., 2016). Under phosphate deficiency, *FRO2* and *IRT1* transcription is strongly downregulated and unlikely to play a major role in Fe reduction and acquisition, at least in the root meristem (Misson et al., 2005; Müller et al., 2007; Thibaud et al., 2010; Lan et al., 2012). Furthermore, the primary roots of both the *fro2* and *irt1* mutants show WT-like responses to Pi deficiency, as shown in this work and that of (Müller et al., 2015), indicating that neither participate in this pathway. Although apoplastic ascorbic acid has been shown to participate in Fe^+3^ reduction for Fe import into developing *A. thaliana* seeds, it is unknown whether ascorbic acid contributes to Fe reduction in the roots (Grillet et al., 2014).

Biochemically characterized CYB561s have shown the ability to use ascorbate as the principal electron donor to generate transmembrane electron transport to reduce either apoplastic monodehydroascorbate or various ferric-chelates (Asard et al., 2013). A CYB561 found in the mouse duodenal mucosa cell membrane transfers electrons from cytoplasmic ascorbate to extracellular Fe^+3^-chelates, enabling the generation of Fe^+2^ ions that are then acquired by Fe^2+^ transporters (McKie et al., 2001; McKie et al., 2002; Ganasen et al., 2018). The characterization of some plant CYB561s showed that they also use ascorbate to generate transmembrane electron transfers to monohydroascorbate or Fe^+3^-chelates when expressed in heterologous systems, such as *Pichia pastoris* (Berczi et al., 2007; Nakanishi et al., 2009a; Nakanishi et al., 2009b). Despite these insights, the *in viv*o relevance of CYB561 proteins to either Fe^+3^ reduction and iron homeostasis or ascorbate pools has not been demonstrated in plants.

The DOMON domain is commonly present as an extracellular domain in membrane-anchored proteins and is often fused to domains other than CYB561 that are involved in redox reactions, such as monooxygenase, cytochrome c, or FAD/NAD-binding oxidoreductase (Aravind, 2001). While only a single CYBDOM protein is found in human, mouse, and *Drosophila*, the CYBDOM protein family is greatly expanded in plants. The DOMON domain of several proteins has been found to bind heme in bacteria and fungi (Hallberg et al., 2000; Kloer et al., 2006). In plants, the AIR12 protein in soybean (*Glycine max*) and *A. thaliana* has a single DOMON domain with a GPI anchor and has been shown to bind heme that is fully reduced by ascorbate (Preger et al., 2009). Although the mode of action of AIR12 is poorly defined, its expression has been associated with superoxide generation at the plasma membrane, the modulation of root development, cold tolerance, and the resistance to *Botrytis cinerea* (Costa et al., 2015; Gibson and Todd, 2015; Biniek et al., 2017; Wang et al., 2021). Studies on a distinct PM-localized CYBDOM from soybean expressed in *Xenopus* oocytes revealed a trans-plasma membrane electron flow dependent on both internal ascorbate and external electron acceptors such as FeCN or ferric nitrilotriacetate (Picco et al., 2015).

The current work shows that CRR is a CYBDOM protein possessing ascorbate-mediated Fe^+3^ reductase activity. CRR contains not only the conserved histidines involved in heme biding in the CYB561 domain, but also the strongly conserved histidine and methionine residues involved in heme *b* binding in the DOMON domain of several bacterial and fungal proteins and the plant AIR12 (Preger et al., 2009; Asard et al., 2013). It is likely that the addition of an extracellular DOMON domain to a CYB561 protein would enable it to transfer electrons to apoplastic acceptors, such as Fe^+3^-chelates, located at a greater distance from the plasma membrane than would be possible with a classical CYB561. Alternatively, the DOMON domain could enable specific interactions with other substrates that will be reduced.

The *crr* mutant shares several phenotypes with the *pdr2*, *als3*, and *star1* mutants, including an enhanced reduction of primary root growth under –Pi, an increased Fe deposition in the apoplast of the SCN, and a reduction in meristem cell division (Ticconi et al., 2009; Müller et al., 2015; Dong et al., 2017). In contrast with *pdr2*, *als3*, and *star1*, *crr* does not affect cell growth in the elongation zone, likely because the accumulation of apoplastic Fe in the elongation zone of *crr* is below a critical threshold. Considering the epistatic interaction between CRR and the *LPR1*, *PDR2*, and *ALMT1* genes, we propose the following model for the implication of CRR on root growth and Fe homeostasis (Figure 6E left). Under P-limiting conditions, the expression of the malate transporter ALMT1 in the roots is induced as the result of STOP1 activation, leading to malate export to the root apoplast (Balzergue et al., 2017; Mora-Macias et al., 2017). This, together with a pool of Fe^+3^ generated by the activity of the ferroxidase LPR1 and the reduced complexing of Fe with Pi, promotes the accumulation of high levels of Fe^+3^-malate in the root apoplast. Fe^+3^-malate reduces division in the root apical meristem through potentially synergic pathways, including the generation of ROS, the activation of the CLE14/CLV2–PEPR2 pathway, and the interference with cell-to-cell communication by callose deposition (Müller et al., 2015; Gutierrez-Alanis et al., 2017). Apoplastic Fe^+3^-malate also reduces cell growth in the root elongation zone, potentially via the activation of peroxidases, cytochrome p450s, and other redox-active enzymes, leading to the modification of the cell wall structure. Blue light is an important component of this process because it activates HY5 in the shoots, promoting its migration to the roots, where it regulates the expression of *LPR1* (Gao et al., 2021). Additionally, blue light triggers photo-Fenton reactions, further inhibiting primary root growth upon their exposure to light (Zheng et al., 2019). In our model, the ascorbate-dependent ferric reductase activity of CRR antagonizes the activity of LPR1, modulating the ratio between the ferric and ferrous iron pools (Fe^+3^/Fe^+2^) in the root apoplast. In the *crr* mutant, the ratio of Fe^+3^/Fe^+2^ would be shifted toward Fe^+3^, increasing the accumulation of Fe^+3^-malate complexes in the apoplast and thereby inhibiting root growth. On the other hand, when *CRR* is expressed in the SCN, the pool of apoplastic Fe^+3^ would decrease via ferric reduction, allowing Fe^+2^ to be transported away from the apoplastic space and into cells, lessening the inhibition of primary root growth under –Pi. CRR protein levels are partially dependent on the ER chaperones CNXs and PDR2 for proper protein expression, folding, and/or localization.

The ferric reductase activity of CRR would also explain the effect of *CRR* expression on a plant’s tolerance or sensitivity to high external Fe (Figure 6E right). In the *crr* and *crr hyp1* mutants, the reduction of ferric reductase activity in the presence of high external Fe would limit the transport of Fe into cells, which would in turn reduce the long-distance Fe transport from root to shoot, preventing its intracellular accumulation to cytotoxic levels in sensitive tissues. Further studies will be required to reveal how the expression patterns of *CRR* and *HYP1* in shoots and germinating seeds contribute to their beneficial effect on germination and seedling establishment under high external Fe. Conversely, in plants overexpressing *CRR*, the increased reduction of Fe^+3^ to Fe^+2^ in the apoplast would lead to a greater Fe^+2^ transport into the roots and transfer to shoots, leading to cytotoxic levels of Fe and the production of ROS, negatively impacting plant fitness.

In most organisms, Fe^+3^ reduction is coupled to Fe^+2^ import either directly via Fe^+2^ transporters, such as IRT1 in plants, which form a complex with FRO2 and the proton pump AHA2 at the PM (Martin-Barranco et al., 2020), or via a coupled ferroxidase/permease transport system, such as the Fet3–Ftr1 high-affinity Fe uptake system of fungi (Stearman et al., 1996; Kosman, 2010). The current work raises the interesting question of how the Fe^+2^ generated by CRR could be coupled to Fe transport. Considering that the *irt1* mutant behaves like the WT as far as the inhibition of primary root growth under –Pi and apoplastic Fe deposition, it is unlikely to be involved in CRR-mediated Fe^+2^ transport, at least in the root meristems (Müller et al., 2015).

Considering the numerous interactions between Fe and Pi homeostasis, it is interesting to highlight that a genetic screen initially devised to study the response of roots to Pi deficiency has led to the identification of key genes likely to be primarily involved in Fe homeostasis, such as CRR and LPR1. Recent studies of the *LPR* gene family in both *A. thaliana* and rice have highlighted their essential contributions to Fe transport and Fe homeostasis in roots and shoots (Ai et al., 2020; Liu et al., 2022; Zhu et al., 2022). Future research on CRR, HYP1, and other members of the CYBDOM and CYB561 protein families should reveal their potential contribution to Fe homeostasis in various tissues.

## Methods

### Plant lines and growing conditions

All Arabidopsis (*Arabidopsis thaliana*) lines and mutants used in this work were in the Columbia (Col-0) background. The mutants and transgenic lines *pdr2-1* (Ticconi et al., 2004), *fro2* (Robinson et al., 1999), *irt1* (Eide et al., 1996), 39OE (Naranjo-Arcos et al., 2017), and pCYCB1::GUS (Colon-Carmona et al., 1999) were previously described. The *hyp1* mutant (At5g35735) was provided by Ricardo Fabiano Giehl (IPK, Gatersleben, Germany). The T-DNA insertion lines *crr-1* (*Salk_202530*), *crr-2* (*Salkseq_039879.2*), *crr-h1* (*Salk_016284, Salk_013527c*), *almt1* (*Salk_009629*) and *lpr1* (*Salk_016297*) were obtained from the European Arabidopsis Stock Center (Nottingham, UK) and genotyped using the primers listed in Supplementary Table 5. All the other genetic constructs used in this work were generated using Infusion and Gateway technology using the primers described in Supplementary Table 5, with the pENTR2b (Thermo Fisher Scientific, Waltham, MA, USA) and pFAST vector series (Shimada et al., 2010) as entry and destination vectors, respectively. Transgenic lines were generated via *Agrobacterium tumefaciens*-mediated transformations (Clough and Bent, 1998).

The seeds were germinated on vertical square plates containing one sixth–strength Murashige and Skoog (MS) salts without Pi (Duchefa Biochemie, Haarlem, the Netherlands), 1% (w/v) sucrose, 0.8% (w/v) agar, 0.5 g l^−1^ of 2-(N-morpholino)ethanesulfonic acid, and buffered to pH 5.8 with KOH. For the +Pi plates, the medium was supplemented with potassium phosphate buffer to obtain a final Pi concentration of 1 mM. For the Fe tolerance assays on plates, the +Pi or –Pi medium was supplemented with FeNaEDTA to the final concentration specified. To assess Fe tolerance in soil, the seeds were sown in soil or clay pellets and watered with tap water or 1/6-strength MS, respectively. Fe was supplemented as FeNaEDTA to the concentration indicated in the text. The plant phenotypes were analyzed after 15 days.

### Proteomic analysis

Col-0 and *cnx1 cnx2* mutant plants were grown in +Pi and –Pi media for seven days before their roots were harvested for a proteomic analysis. The tissues were homogenized in SDS-containing FASP buffer (4% SDS, 0.1 M DTT, 100 mM Tris pH 7.5), heated at 70°C for 5 min, and allowed to cool at room temperature (RT). The cooled proteins were centrifuged for 20 min at 4000 rpm, after which the supernatant was recovered and frozen at –80°C. The concentration of each extract was determined by gel electrophoresis, Coomassie blue staining, and total lane densitometry in comparison with a pre-quantitated total cell extract, and 50 µg of each tissue extract was subjected to further processing. Four replicates for each condition were prepared. The eight-plex iTRAQ approach (Hunt et al., 2004), based on isobaric labeling and the measurement of reporter fragment ion intensities, was used for the relative quantitation. The extracts were proteolytically digested and iTRAQ-labeled according to the iFASP procedure, which was performed essentially as previously described (McDowell et al., 2013). More than 98.5% of the sequences of all samples had peptide spectrum matches determined by MASCOT (www.matrixscience.com) that corresponded to fully labeled iTRAQ peptides.

The labeled iTRAQ samples (8 × 50 µg) were pooled and desalted on SepPak C18 cartridges (Waters Corporation, Milford, MA, USA). Dried eluates were dissolved in 4 M urea with 0.1% ampholytes (pH 3–10; GE Healthcare, Chicago, IL, USA) and fractionated by off-gel focusing, as previously described (Geiser et al., 2011). The 24 fractions obtained were desalted on a microC-18 96-well plate (Waters Corporation), dried, and resuspended in 0.1% formic acid with 3% (v/v) acetonitrile for LC-MS/MS analysis.

A data-dependent LC-MS/MS analysis was performed on the extracted peptide mixtures after digestion using a Fusion tri-hybrid orbitrap mass spectrometer (Thermo Fisher Scientific) interfaced through a nano-electrospray ion source to a RSLC 3000 HPLC system (Thermo Fisher Scientific). The peptides were separated on a reversed-phase Easy-spray PepMap nanocolumn (75 μM inner diameter × 50 cm, 2 μM particle size, 100 Å pore size; Thermo Fisher Scientific) with a 4–76% acetonitrile gradient in 0.1% formic acid (total time: 140 min). Full MS survey scans were performed at 120’000 resolution. In the data-dependent acquisition controlled by Xcalibur 3.0.63 software (Thermo Fisher Scientific), the most intense multiply charged precursor ions detected in the full MS survey scan were selected for “multi-notch” synchronous precursor selection (McAlister et al., 2014) and MS3 fragmentation, which minimizes the co-isolation of near-isobaric precursors (maximum scan cycle: 3 s).

The data files were analyzed with MaxQuant 1.5.3.30 (Cox and Mann, 2008) incorporating the Andromeda search engine (Cox et al., 2011). The sequence database used for the search was the UNIPROT *Arabidopsis thaliana* proteome containing 27,242 sequences, in addition to a custom database containing most of the common environmental contaminants (keratins, trypsin, etc.). Both peptide and protein identifications were filtered at a 1% FDR relative to hits against a decoy database built by reversing the protein sequences. The raw reporter ion intensities generated by MaxQuant were used in all following steps leading to the quantification. Further data analysis was performed using the R statistical programming language version 2.15.2 (R Core Team, 2012) and the Perseus software (Cox and Mann, 2008).

### Root length quantification

The plants were grown as described above and the plates were imaged with a flatbed scanner (Epson Perfection V700 photo; Epson, Suwa, Japan). The root length was measured using Fiji (http://fiji.sc/Fiji) and the plugin Simple neurite tracer.

### Determination of meristem size and cell length

The roots were cleared with Clearsee and stained with Calcofluor White, as previously described (Ursache et al., 2018), and observed using confocal microscopy. The average cell length of the cortical cells in the differentiation zone was calculated. The root apical meristem size was determined as the distance from the QC to the first elongating cell.

### Confocal microscopy

Confocal microscopy was performed using a Zeiss LSM 880 Airyscan (Carl Zeiss). Calcofluor White was excited at 405 nm and detected at 425–475 nm. GFP was excited with a 488 nm laser and the emitted light was collected at 493–538 nm.

### Measurement of Pi content

Pi was quantified using the ascorbate–molybdate assay, as previously described (Ames, 1966). Around 20 mg of shoot or root tissue was placed in water and subjected to at least three freeze-thaw cycles to release the Pi. The Pi context was quantified using a molybdate assay with a standard curve. The Pi concentration was normalized per mg of tissue.

### Beta-glucuronidase (GUS) assay

Col-0 and *crr-1* expressing the *pCYCB1::GUS* reporter cassette grown in +Pi and –Pi conditions were incubated in the GUS staining buffer (50 mM sodium phosphate, 2 mM potassium ferrocyanide, 2 mM potassium ferricyanide, 0.2% Triton X-100, and 2 mM of the GUS substrate 5-bromo-4-chloro-3-indolyl-b-D-glucuronic acid), as previously described (Jefferson et al., 1987). The roots were imaged using a bright field stereomicroscope (Axio Zoom.V16; Carl Zeiss, Oberkochen, Germany).

### Measurement of ferric-chelate reductase activity in roots

Roots from seven-day-old seedlings grown on 1/6-strength MS plus Pi were used to measure Fe(III) chelate reductase activity. The root tissue was washed with a 100 mM Ca(NO_3_)_2_ solution and incubated for 1 h in the dark with 1 ml of reaction solution. For the Fe(III)NaEDTA reduction, a 100-µM FeNaEDTA and 500-µM ferrozine solution was added to each sample, and the production of purple-colored Fe(II)–ferrozine complexes was measured by measuring the absorbance of the solution at 562 nm (Naranjo-Arcos et al., 2017). For the ferricyanide (FeCN) reduction, the reaction solution was 500 µM FeCN, and the reduction of ferricyanide was recorded by a spectrophotometer at 420 nm (Doring and Luthje, 2001).

### Measurement of ferric-chelate reductase activity in yeast

*Saccharomyces cerevisiae* strain S288C Wt or *Δfre1Δfre2* cells (Georgatsou and Alexandraki, 1994) were used for the ferric-chelate reductase activity assay. For this purpose, the cells were transformed with the empty centromeric p415 vector (Mumberg et al., 1995) as a negative control, or the same vector carrying the *CRR* or yeast *FRE1* coding sequence under the control of the constitutive *GPD* promoter. For the assay, the cells were grown on solid SD/-LEU media for two days and then overnight in 5 ml liquid media with or without an ascorbate precursor (89 mg/100 ml L-galacto-γ-lactone or 25 mg/100 ml L-galactose) until a saturated culture was obtained (Spickett et al., 2000; Berczi et al., 2007). The cells were washed twice with H_2_O, and 1 ml of the resuspension was placed in a 2 ml tube for the analysis. The cells were collected by centrifugation and resuspended in 650 µl of assay buffer consisting of 0.05 M sodium citrate buffer (pH 6.5) and 5% glucose supplemented with 500 µM ferrozine and 100 µM FeNaEDTA or 500 µM FeCN. After a 4-h incubation, an aliquot was collected to record the optical density at 600 nm and the rest was centrifuged to pellet the cells. The optical density of the supernatant was recorded at 562 nm or 420 nm for the FeNaEDTA or FeCN reduction, respectively.

### Histochemical detection of Fe in seedlings

Fe accumulation in seedlings was assayed using Perls or Turnbull staining as indicated (Meguro et al., 2007). For Perls staining, the seedlings were incubated in 4 ml of 2% (v/v) HCl and 2% (w/v) potassium ferrocyanide for 30 min. The samples were washed with H_2_O and incubated for 45 min with 4 ml of 10 mM NaN_3_ and 0.3% H_2_O_2_ in methanol. After this step, the samples were washed with 100 mM Na-phosphate buffer (pH 7.4) and were either analyzed or subjected to a DAB intensification step. For the latter, the seedlings were incubated for 30 min in the same buffer containing 0.025% (w/v) DAB and 0.005% (v/v) H_2_O_2_ and washed two times with H_2_O. Finally, the samples were cleared with chloral hydrate (1 g ml^−1^; 15% glycerol) and analyzed with an optical microscope. The Turnbull method was carried out in the same way, but replacing the potassium ferrocyanide with potassium ferricyanide.

### ^55^Fe transport in Arabidopsis tissues

Plants were grown for seven days in plates containing 1/6-strength MS supplemented with 0.8% (w/v) of agar and transferred to plates supplemented with 600 µM FeNaEDTA and 1.67 µCi ml^−1^ of ferric ^55^Fe for four days. After this, the root and shoot tissues were harvested separately and washed with cold 1/6-strength MS. The samples were dried for two days at 60°C before being digested with 0.2 ml perchloric acid at 90°C for 8 h. The solution was cleared by adding 0.4 ml H_2_O_2_ and incubating overnight at 90°C. For the ^55^Fe quantification, 4 ml of HionicFluor scintillating mixture (PerkinElmer, Waltham, MA, USA) was added to the samples, and the counts per minute were measured with a Tri-Carb 2800TR liquid scintillation counter (PerkinElmer) (Grillet et al., 2014).

### ROS quantification

ROS quantification was performed according to a published protocol (Bellegarde et al., 2019). Briefly, approximately 40 mg of tissue was ground in liquid nitrogen and resuspended in 400 µl of phosphate buffer. The solution was cleared by centrifugation at 13,000 × g for 5 min at 4°C, and an aliquot of 10 µl was mixed with 190 µl of carboxy-H2DCFDA in PBS (to a final concentration of 25 µM carboxy-H2DCFDA) and incubated for 30 min. The florescence emission was quantified as relative fluorescence units.

### RNA extraction and qRT-PCR

Total RNA was extracted using the ReliaPrep RNA tissue miniprep system kit (Promega, Madison, WI, USA) and treated with RNase-free DNase according to the manufacturer’s instructions. The first-strand cDNA was synthetized using M-MLV reverse transcriptase (Promega). The qRT-PCR was performed in a Quantstudio 3 thermocycler (Thermo Fisher Scientific) using the SYBR select master mix (Thermo Fisher Scientific) and the primers listed in Supplementary Table 5. *ACTIN2* (At3g18780) was used as a reference gene to normalize the gene expression data. Three biological replicates and two technical replicates were performed for each gene.

### RNA-seq analysis

Col-0 and CRR OE were grown in +Pi and –Pi conditions for seven days, as described above, and root tissues from 60 seedlings were pooled in each biological replicate, from which total RNA was extracted using Trizol (Thermo Fisher Scientific). RNA-seq was performed on three biological replicates per condition using Illumina (San Diego, CA, USA) technology, with more than 25 million reads obtained for each of them. Adapters were removed from the purity-filtered reads, which were trimmed for quality using Cutadapt v. 1.8 (Martin, 2011). Reads matching ribosomal RNA sequences were removed using fastq_screen v. 0.11.1 (https://www.bioinformatics.babraham.ac.uk/projects/fastq_screen/). The remaining reads were further filtered for low complexity with reaper v. 15-065 (Davis et al., 2013) and aligned against the *Arabidopsis thaliana* TAIR10.39 genome using STAR v. 2.5.3a (Dobin et al., 2013). The number of read counts per gene locus was summarized with htseq-count v. 0.9.1 (Anders et al., 2015) using the *Arabidopsis thaliana* TAIR10.39 gene annotations. The quality of the RNA-seq data alignment was assessed using RSeQC v. 2.3.7 (Wang et al., 2012). Genes with low counts were filtered out according to the rule of 1 count per million in at least one sample. The library sizes were scaled using TMM normalization and log-transformed into counts per million using the EdgeR package version 3.24.3 (Robinson et al., 2010). Differential expression levels were computed with limma-trend approach (Ritchie et al., 2015) by fitting all samples into one linear model. A moderated t-test was used for each contrast and an F-test was used for the interaction. For each group of comparisons, the adjusted p-value is computed by the Benjamini–Hochberg method, controlling for the false discovery rate (FDR or adj.P.Val).

## Gene editing using CRISPR/Cas9

The vectors used to perform the CRISPR/Cas9-mediated *CRR* gene editing were previously described (Ursache et al., 2021). Briefly, sgRNAs were designed using CRISPR-P V2 (http://crispr.hzau.edu.cn/CRISPR2/) and cloned into the entry vectors pRUs. Three different sgRNAs were combined into the pSF278 intermediate vector, and finally recombined into the T-DNA destination vector PcUBi4-2::SpCas9 FastRed.

## Accession numbers

The Arabidopsis accession numbers used in this work are the following: CRR, AT3G25290; CRR-H1, AT4G12980; HYP1, AT5G35735; PDR2, AT5G23630; LPR1, AT1G23010; ALTM1, AT1G08430; BHLH39, AT3G56980; FRO2, AT1G01580; IRT1, AT4G19690; CNX1, AT5G61790; CNX2, At5G07340.

The RNA seq dataset has been deposited at the Gene Expression Omnibus (GSE215755, token for reviewers gfunmymkltctzyv).

## Acknowledgments

The authors acknowledge Manfredo Quadroni and Patrice Waridel of the Protein Analysis Facility and the team in the Genomic Technology Facility, University of Lausanne, Switzerland, for the proteomic and transcriptomic analyses, respectively. The support of the University of Lausanne Cellular Imaging Facility is also acknowledged. The authors are grateful to Ricardo Fabiano Giehl and Nicolaus von Wiren (IPK, Gatersleben, Germany) for providing seeds of the *hyp1* mutant, Steffen Abel (Leibniz Institute, Halle, Germany), Thierry Desnos (CEA, Cadarache, France) and Petra Bauer (Heinrich Heine Universität, Düsselorf, Germany) for sharing seeds of the *pdr2*, *lpr1*, and *almt* mutants and 39 OE transgenic line, and Diane Ward (University of Utah, Salt Lake City, USA) for the yeast mutant Δ*fre1 fre2*. The authors are also grateful to Christian Dubos and Christian Hardtke for helpful discussions. This work was supported by a grant from the Swiss National Science Foundation (31003A-182462) to YP.

## Author contributions

JC and YP conceived the project. JC performed all experiments, with the exception of the root proteomics performed by JM. JC and YP wrote the manuscript, JM provided feedback on it, and all authors approved the final manuscript. YP agrees to serve as the author responsible for communication.

