## Supplemental Figures for "A DOMON domain-cytochrome b561 protein acts as a ferric reductase in iron homeostasis and impacts primary root growth under phosphate deficiency"

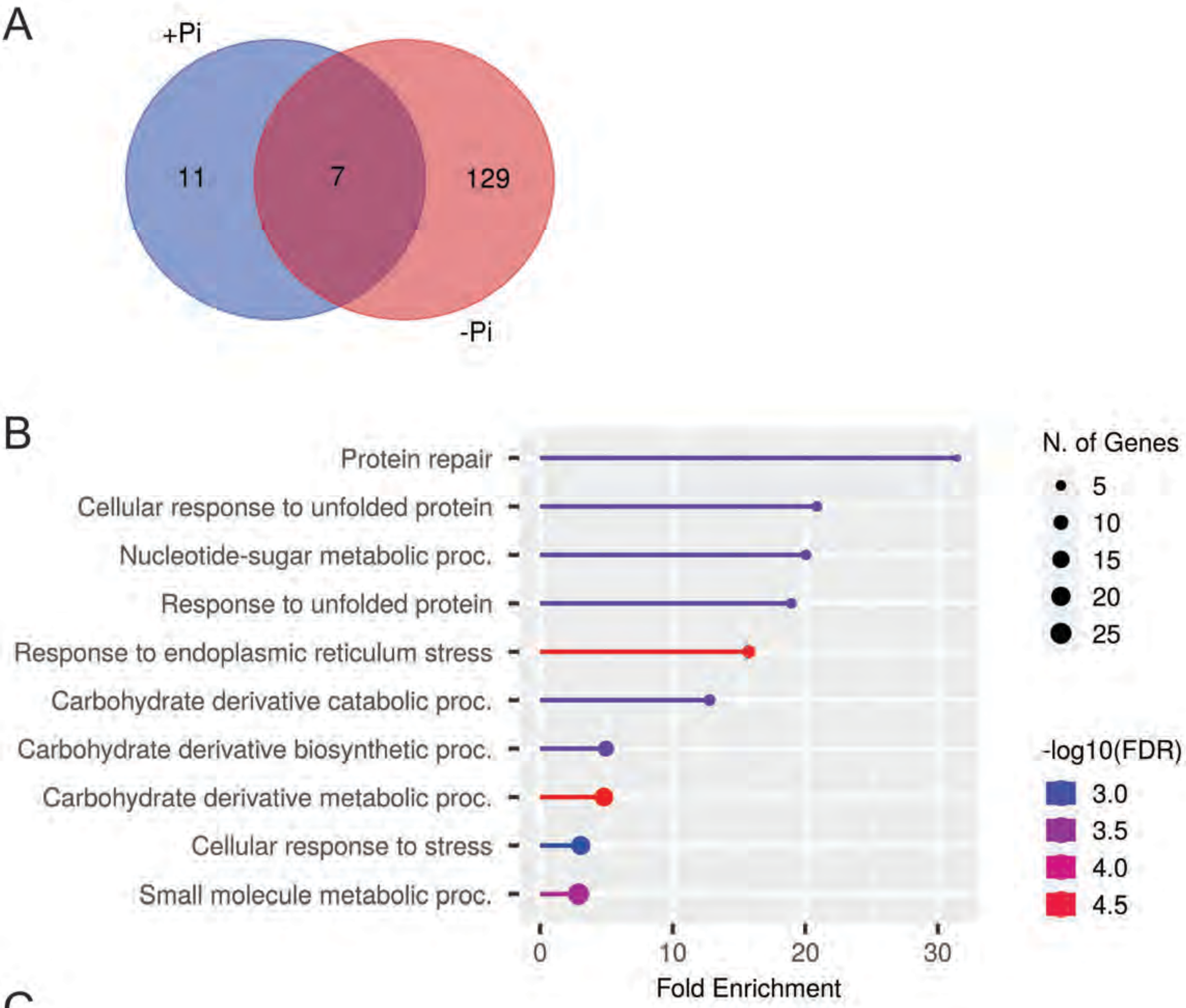


C

| | |
|---|---|
| AT3G45600 | Member of TETRASPANIN family (TET3) |
| AT3G25290 | Cytochrome and domon domain containing protein |
| AT1G76090 | Sterol methyltransferase 3 (SMT3) |
| AT2G28190 | Copper zinc superoxide dismutase 2 (CSD2) |
| AT4G16660 | Heat shock protein 70 (Hsp 70) family protein |
| AT5G13640 | Phospholipid:diacylglycerol acyltransferase (PDAT) |
| AT1G04910 | O-fucosyltransferase family protein |

**Figure S1. Proteomic analysis of *cnx1 cnx2* double mutant.**
**A**, Venn diagram showing the differentially abundant proteins between Col-0 and *cnx1 cnx2* in plus (+Pi) and minus (-Pi) phosphate conditions. Plants were grown for 7 days in +Pi and -Pi and root tissue was processed for a proteomic analysis based on iTRAQ labeling and LC-MS/MS. For each condition differential abundant proteins between Col-0 and *cnx1 cnx2* were identified according to the following criteria: p-value ≤0.05 Student's T-test; -1.5 ≥ FC ≥ 1.5. **B**, Gene ontology (GO) enrichment analysis of the differentially abundant proteins between Col-0 and *cnx1 cnx2*. The top 10 enriched 'molecular function' GO terms are shown in a lollipop graph where the fold enrichment, number of the corresponding genes (N. of genes) and the fold discovery rate (FDR) are indicated.
**C**, Table showing the seven proteins that were differentially abundant between Col-0 and *cnx1 cnx*2 both in +P and -Pi. The gene identification number and the annotation is shown.

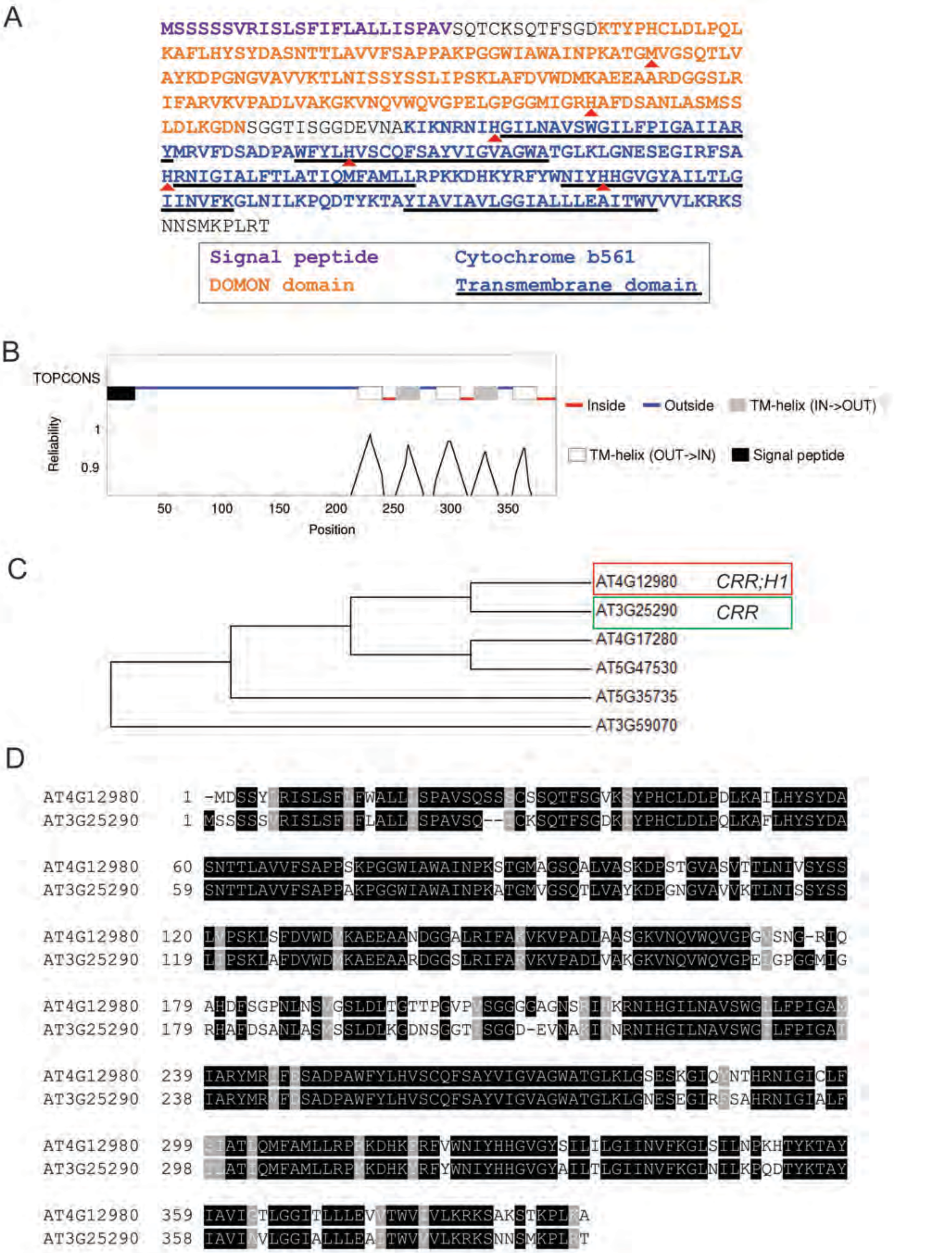


**Figure S2. CRR protein features and phylogeny**
**A**, CRR amino acid sequence showing the location of the signal peptide (SP), the DOMON, cytochrome b561 and transmembrane (TM) domains. The conserved methionine (M) and histidine (H) of the DOMON domain as well as the four conserved histidines of the cytochrome b561 are indicated by red triangles. **B**, CRR topology and transmembrane domains as predicted by the TOPCONS software (topcons.cbr.su.se). **C**, Phylogenetic tree showing the 6 Arabidopsis CYBDOM members belonging to the F2 clade. CRR and CRR-H1 are highlighted in green and red, respectively. The tree was constructed using the neighbor-joining method based on the multiple sequence alignment analysis of the full-length proteins. **D**, Protein sequence alignment of CRR and CRR-H1. The alignment was generated using ClustalOmega and the conserved regions highlighted using BOXSHADE.

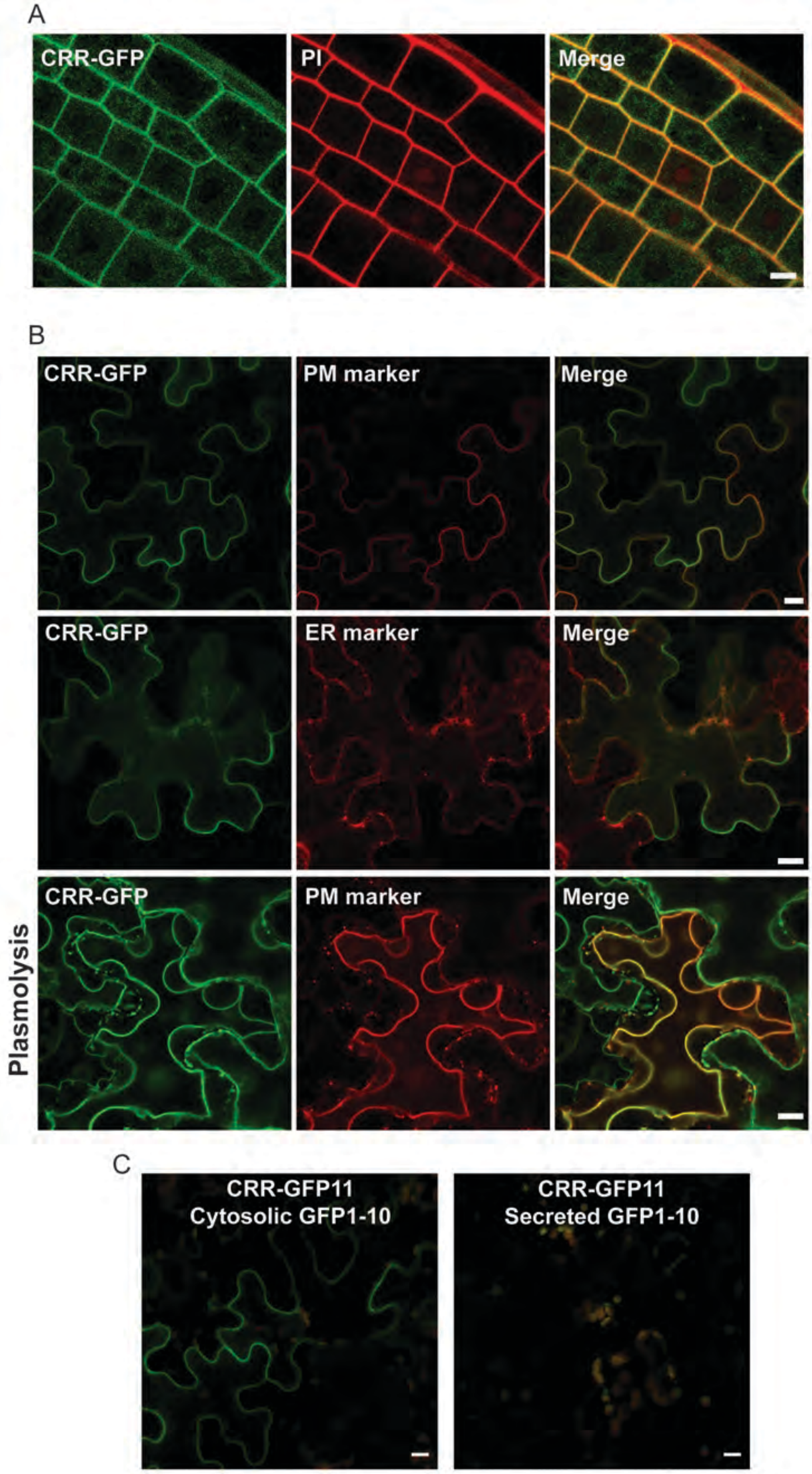


**Figure S3. CRR subcellular localization and topology analysis**
**A**, CRR primarily localizes to the plasma membrane in *Arabidopsis thaliana*. The confocal microscopy images show an Arabidopsis transgenic root expressing a translational fusion CRR-GFP driven by the 35S promotor (green channel) together with the cell wall marker propidium iodide (PI) in red. The merged channels show the co-localization between CRR and PI. **B**, Subcellular localization of CRR in *Nicotiana benthamiana*. Confocal images showing *N. benthamiana* epidermal cells expressing CRR-GFP together with the plasma membrane marker CBL1 (top panels) fused to the orange fluorescent protein or the ER marker ER-rk fused to mCherry (middle panels). In order to confirm CRR plasma membrane localization cells, were plasmolyzed by the addition of NaCl (bottom panels). **C**, Split-GFP system showing that CRR C-terminus is facing the cytosol. The 11th beta-strand of the GFP was translationally fused to the C-terminus of CRR (CRR-GFP11) and co-expressed in *N. benthamiana* together with the other 10 GFP beta-strands targeted either to the cytosol (cytosolic GFP1-10) or the apoplast (secreted GFP1-10). GFP fluorescence was only detected by confocal microscopy when CRR-GFP11 was co-expressed with the cytosolic GFP1-10, indicating that the CRR C-terminus is intracellular. In (A), (B) and (C) scale is 10 µm.

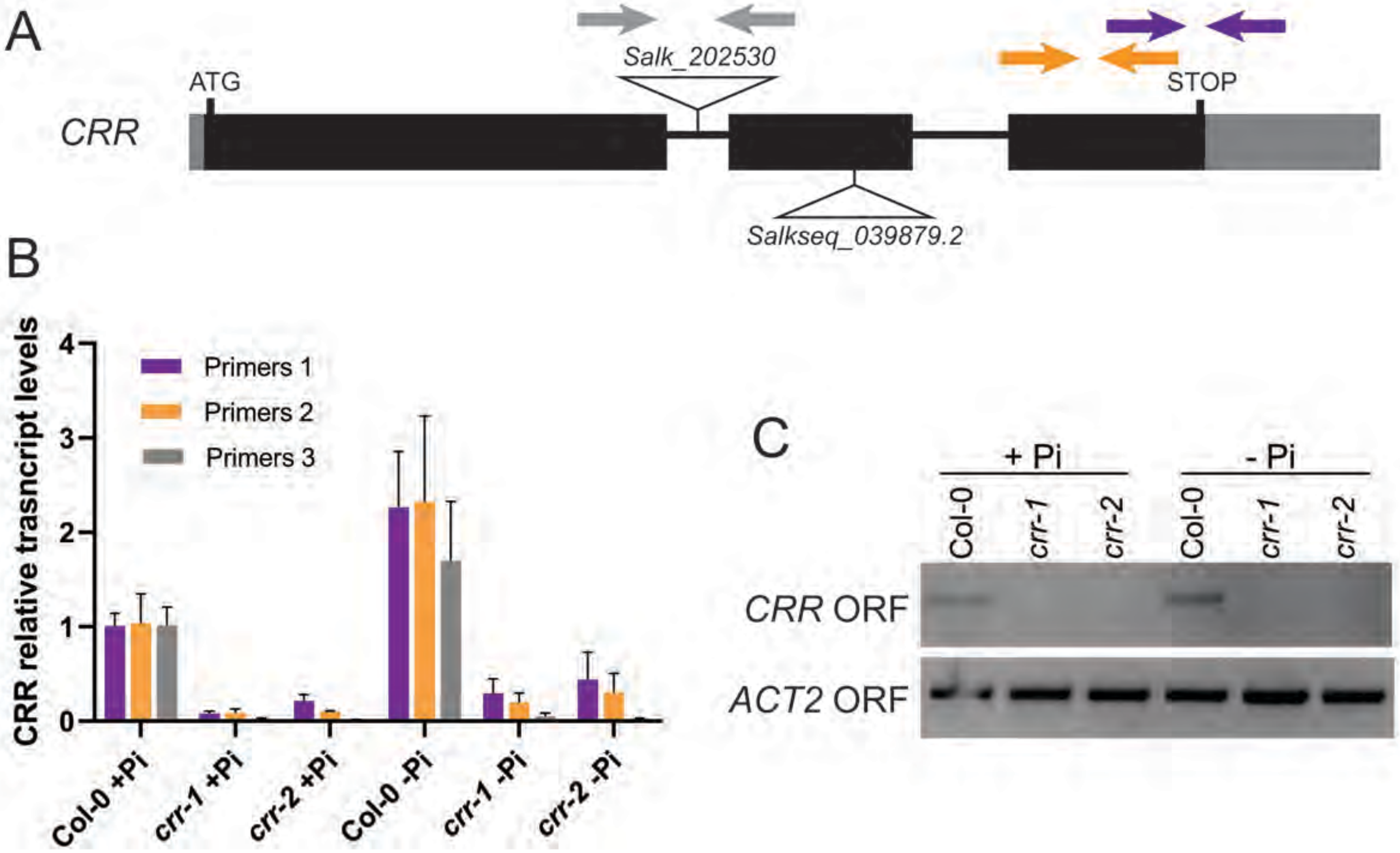


**Figure S4. Characterization of *Arabidopsis thaliana* CRR T-DNA mutants**

**A**, Scheme showing the *CRR* gene structure and the localization of two different T-DNA insertions. Grey boxes indicates the 5'- and 3'-UTRs, black boxes represents the 3 exons and the black lines connecting them the introns. The *A. thaliana* line Salk_202530 (*crr-1*) has a T-DNA insertion in the first intron while the line Salkseq_039879.2 (*crr-2*) has an insertion in the second exon. The grey, orange and purple arrows indicate the positions of the primers used in (B). **B**, The two *CRR* T-DNA mutants *crr-1* and *crr-2* have reduced levels of CRR mRNA. Arabidopsis Col-0, *crr-1* and *crr-2* seedlings were grown for 7 days in plus (+Pi) or minus (-Pi) phosphate conditions and *CRR* relative transcript levels were quantified by qRT-PCR using the three different set of primers shown in (A). **C**, Full length *CRR* complementary DNA (cDNA) is undetectable in the *crr* T-DNA mutants. cDNA obtained from Col-0, *crr-1* and *crr-2* grown in +Pi and -Pi was utilized as template for PCR reactions using primers to amplify either *CRR* or *ACT2* complete open reading frames (ORF). The PCR products were resolved by agarose gel electrophoresis and revealed by ethidium bromide staining. While *ACT2* was detected in all samples, *CRR* ORF was only detected in Col-0 samples.

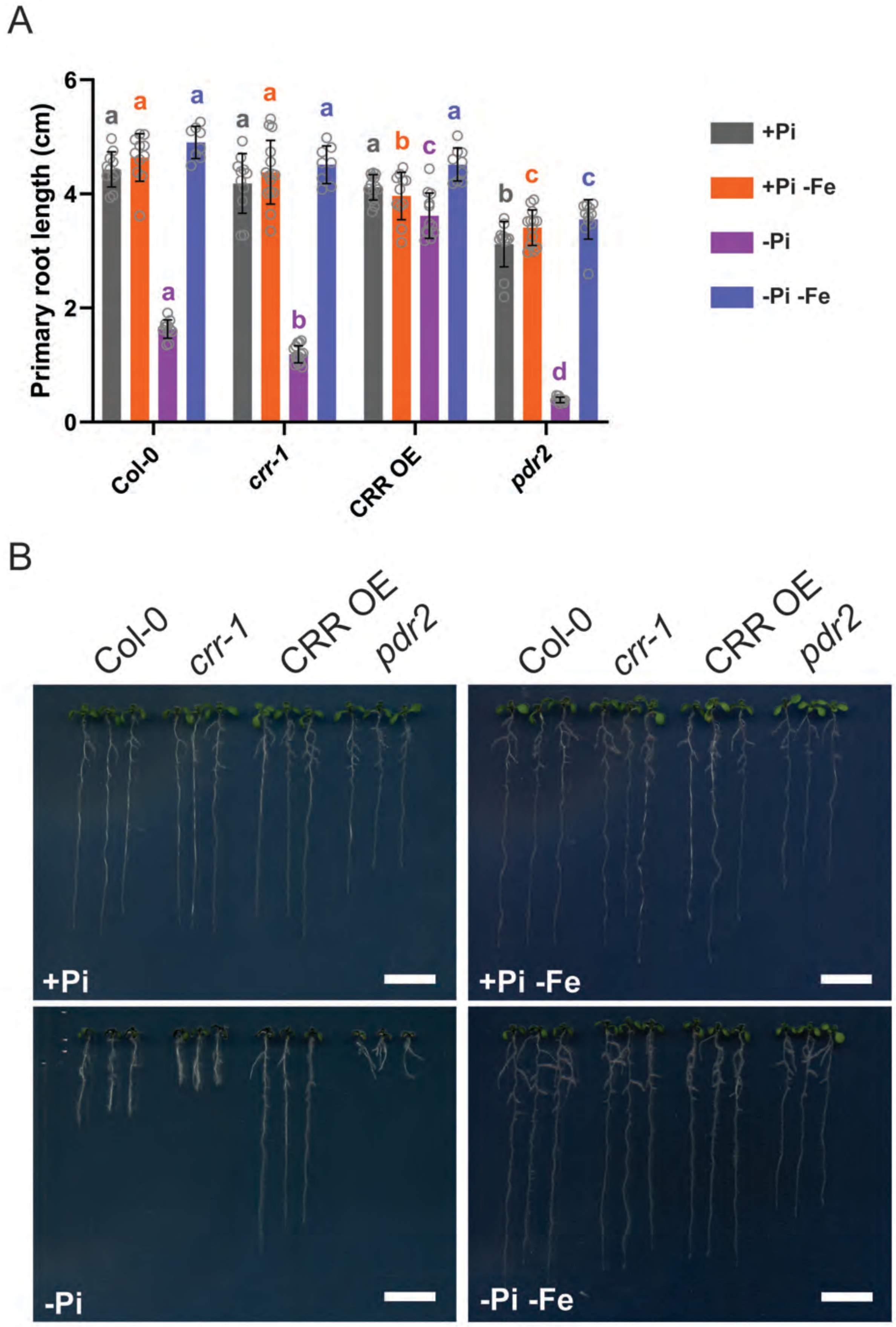


**Figure S5. *crr* primary root growth phenotype under phosphate starvation is iron dependent**
A, Primary root length quantification of Col-0, *crr-1*, CRR OE and *pdr2* seedlings grown in plus (+Pi) or minus phosphate (-Pi) with or without the addition of ferrozine to sequester the iron in the media. Media with the addition of ferrozine is indicated as (-Fe). The data is represented as mean ± s.d. (n≥8) and a Two-way ANOVA followed by a Tukey's multiple comparisons test was used to assess the statistical differences. Significant differences ($P < 0.05$) within each condition (same color code) are indicated by different letters. B, Photographs showing representative 7 days post germination seedlings grown in the conditions indicated in (A). Scale: 1cm.

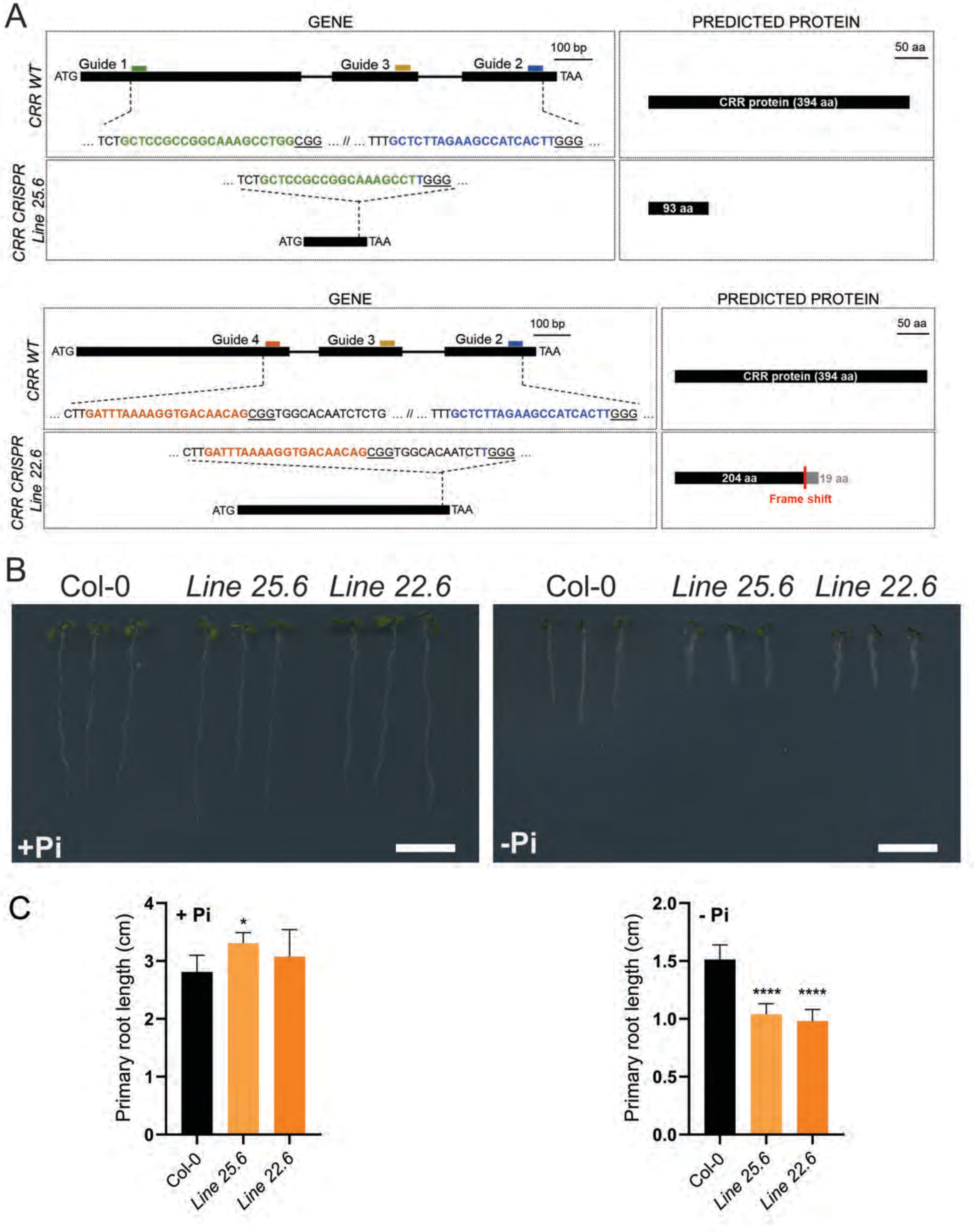


**Figure S6. *crr* mutants generated by CRISPR/Cas9 show a short-root phenotype**
**A**, Schematic representation of the strategy used to edit the *CRR* locus by CRISPR/Cas9 and the resulting edited lines. Two pools of Arabidopsis plants were transformed with different sets of guide RNAs (guides 1, 2, and 3 or guides 2, 3 and 4) targeting the *CRR* locus. The analysis of independent T1 plants allowed the identification of two independent edited lines. Line 25.6 has an in-frame deletion of 1120 bases in the *CRR* gene, generating an open reading frame coding for an hypothetical protein of 93 amino acids (top panel). Line 22.6 bears a frame-shift mutation of 729 nucleotides generating an hypothetical ORF coding for a protein of 223 amino acids where the first 204 corresponds to the DOMON domain (bottom panels). **B**, Pictures showing the root phenotype of the *crr* edited lines grown in plus (+Pi) and (-Pi) minus phosphate conditions. Scale: 1 cm. **C**, Primary root length quantification of Col-0 and the *crr* edited lines 7 days after germination. Data represents the mean ± s.d. (n≥7) and the statistical differences were analyzed using a One-way ANOVA followed by a Tukey's multiple comparisons test. Significant differences against Col-0 are indicated (***P < 0.001; **P < 0.01; *P < 0.05).

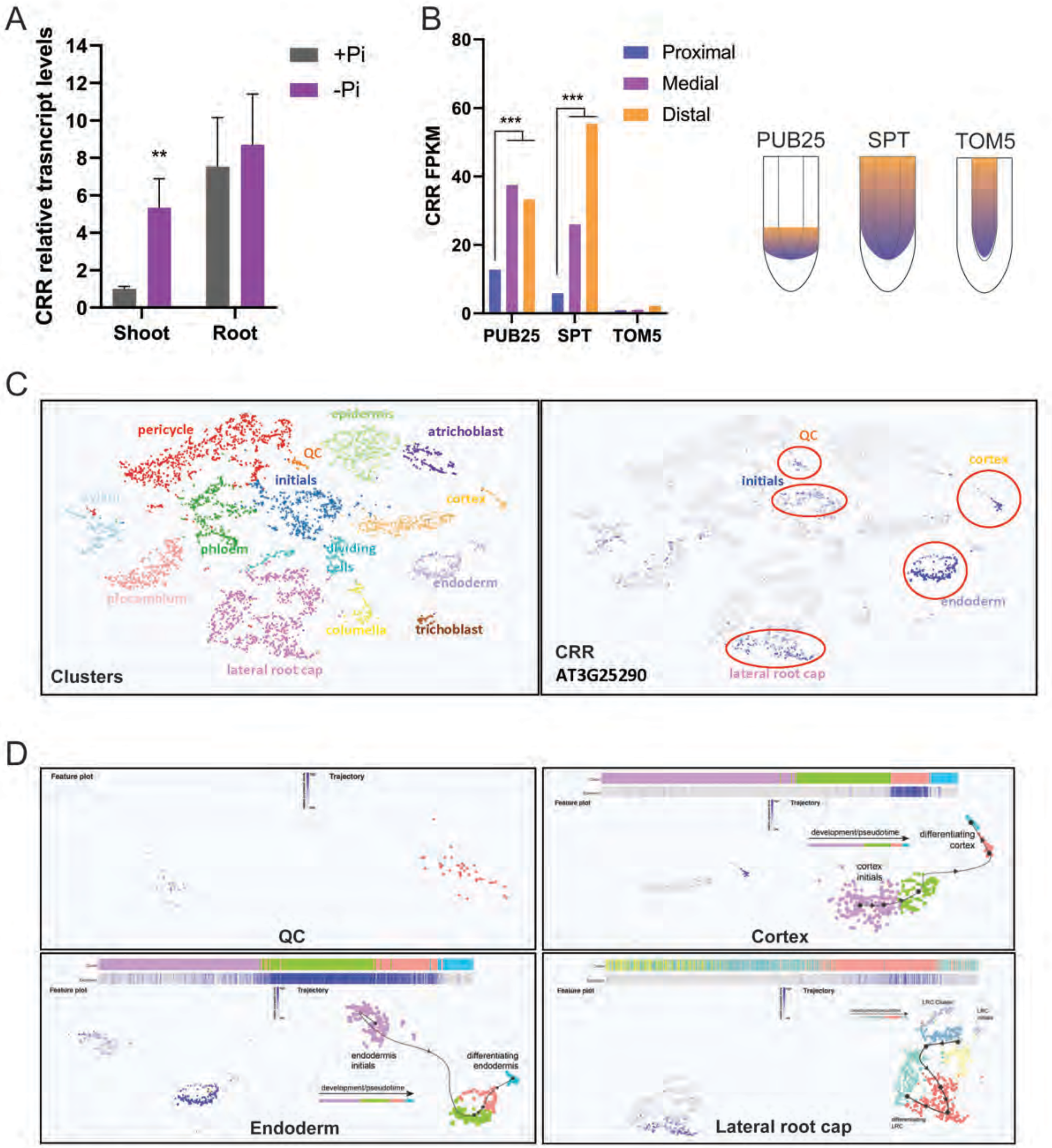


**Figure S7. *CRR* expression pattern in root tips.**
**A**, *CRR* is induced by phosphate starvation in shoots. Graph showing *CRR* relative transcripts levels in shoots and roots of plants grown for 7 days in plus (+Pi) and minus (-Pi) phosphate. Data is represented as the mean of 3 biological replicates ± s.d. For each tissue significant differences between +Pi and -Pi were analyzed using an unpaired t-test (**$P < 0.01$). **B**, *CRR* is expressed in the proximal root apical meristem. Data retrieved from Wendrich et al., 2017 (DOI: 10.1073/pnas.1707400114 ) showing *CRR* expression levels in root cell populations which express the *PUB25*, *SPT* or *TOM5* gene. The scheme indicates the expression pattern of these 3 genes as well as the intensity of the expression along the root. **C**, Single-cell transcriptomic data from Arabidopsis root tips was retrieved from the Plant sc-Atlas (https://bioit3.irc.ugent.be/plant-sc-atlas/). The left panel shows a color-coded tSNE plot with the classification of sequenced cells into the different cell identities, and the right panel shows which cells express *CRR*. **D**, Single-cell transcriptomic data from Arabidopsis roots showing the expression of the *CRR* gene in QC, endodermis, cortex and lateral root cap cells in relation to the developmental trajectories of the same cells.

A

| Promoter name | Id Gene | Cell type | Reference |
|---|---|---|---|
| LPR1 | At1G23010 | Stem cell niche (SCN), endoderme and provascular tissues | Muller et al., 2015. doi: 10.1016/j.devcel.2015.02.007 |
| SCR | At3g54220 | Root endodermis, quiescent center | Mustroph et al., 2009. doi: 10.1073/pnas.0906131106 |
| MYB36 | At5G57620 | Endoderme elongation | Pengxue Li et al., 2018. doi: 10.1016/j.cub.2018.07.028 |
| WOL | At2g01830 | Root vasculature | Mustroph et al., 2009. doi: 10.1073/pnas.0906131106 |
| CO2 | At1g62500 | Root cortex meristematic zone | Mustroph et al., 2009. doi: 10.1073/pnas.0906131106 |
| PEP | At1g09750 | Root cortex elongation and maturation zone | Mustroph et al., 2009. doi: 10.1073/pnas.0906131106 |
| ATL75 | At1G49200 | Pericycle | Denyer et al., 2019. doi: 10.1016/j.devcel.2019.02.022 |
| LOVE1 | AT5G20045 | Root cap | Berhin et al., 2019. doi: 10.1016/j.cell.2019.01.005 |
| WOX5 | At3G11260 | Quiescent center (QC) | Burkart et al., 2022. doi: 10.15252/embr.202154105 |

B

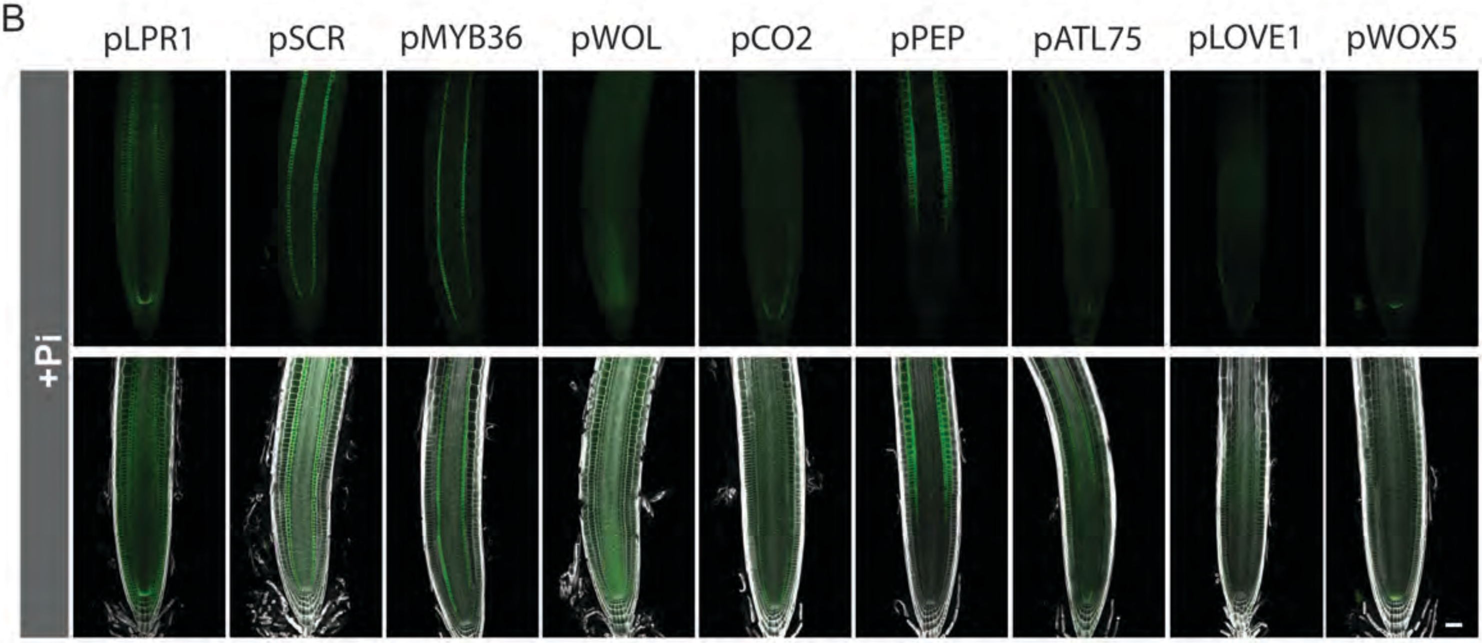


C

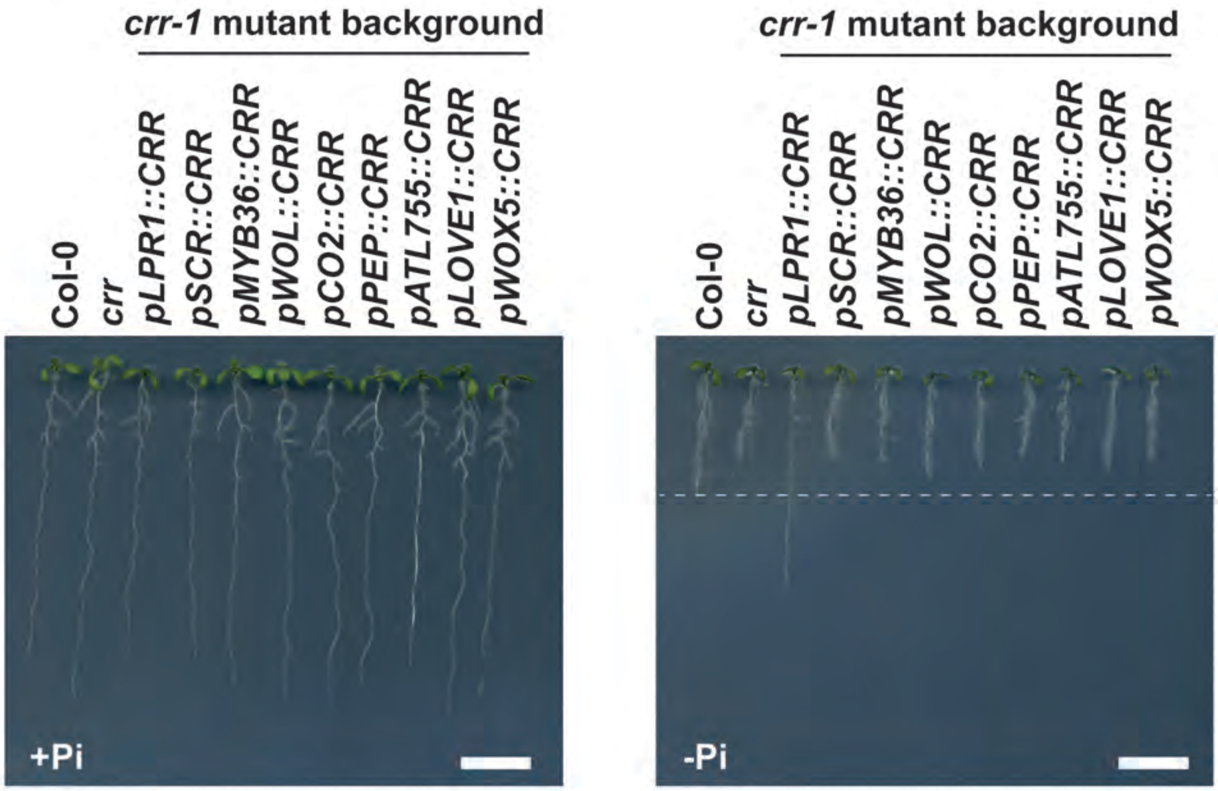


**Figure S8. *CRR* expression driven by *LPR1* promoter complements *crr* mutant phenotype**
A, Description of the promoters used to drive *CRR-GFP* expression. B, Confocal microscopy images of *crr* mutant roots showing the expression of CRR-GFP translational fusion under the control of different promotors. B, Images showing the primary root phenotype of the transgenic lines shown in (A).

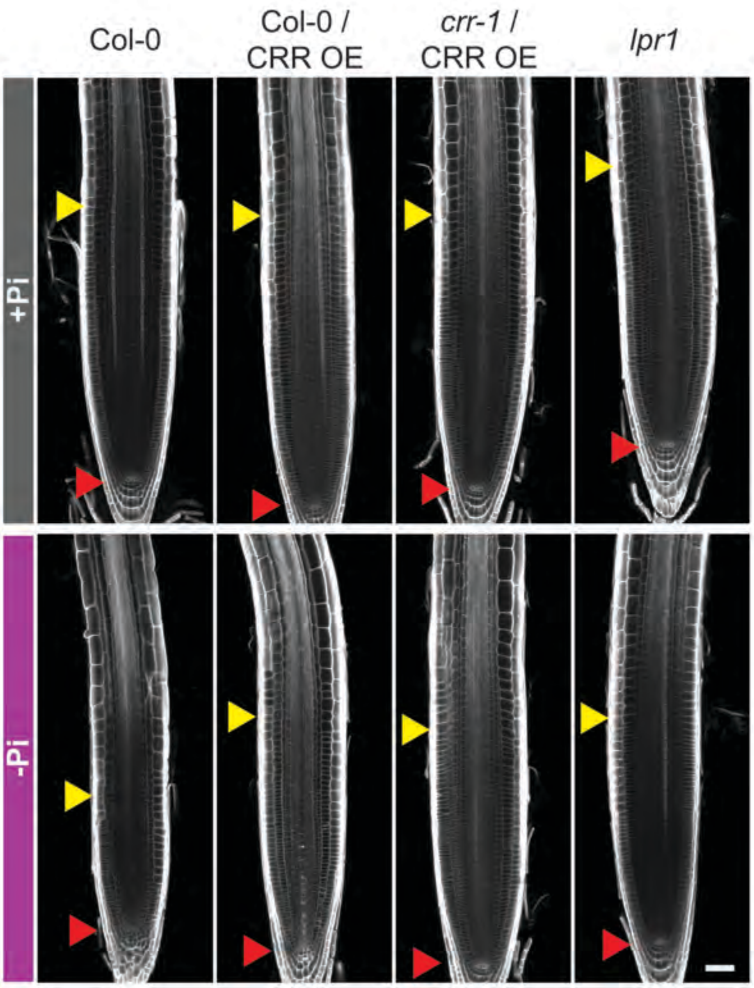


**Figure S9. The root apical meristem of CRR OE lines is insensitive to phosphate starvation.**
Confocal microscopy images showing the size and morphology of the root apical meristem of Col-0, CRR over-expressing lines (CRR OE) in a Col-0 or *crr-1* mutant background and *lpr1* seedlings grown in plus (+Pi) or minus (-Pi) phosphate conditions. Cell walls were stained using Calcofluor white. The quiescent center and the end of the meristem are indicated by red and yellow triangles, respectively. Scale: 20 µm

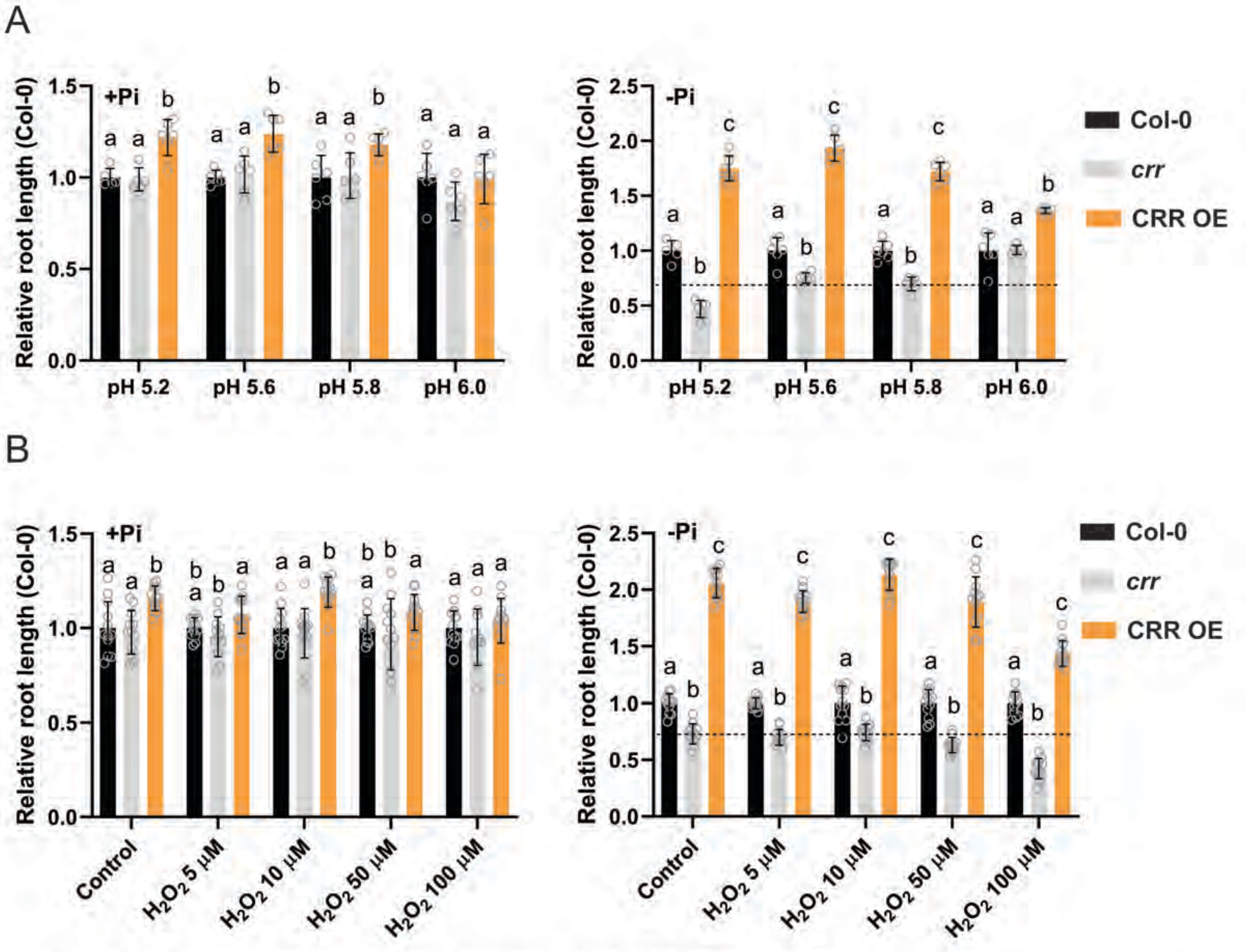


**Figure S10.** ***crr* mutant phenotype is enhanced by low pH and hydrogen peroxide**
A, Primary root length of *crr-1* and CRR OE relative to Col-0 when plants where grown in plus (+Pi) or minus (-Pi) phosphate at different pH. B, Primary root length of *crr-1* and CRR OE relative to Col-0 when seedlings where grown in (+Pi) or (-Pi) phosphate and increasing concentrations of hydrogen peroxide (H2O2). In (A) and (B) the data is represented as mean ± s.d. (n≥5).
A Two-way ANOVA followed by a Tukey's multiple comparisons test was used to assess the statistical differences. Significant differences ($P < 0.05$) within each condition (same color code) are indicated by different letters.

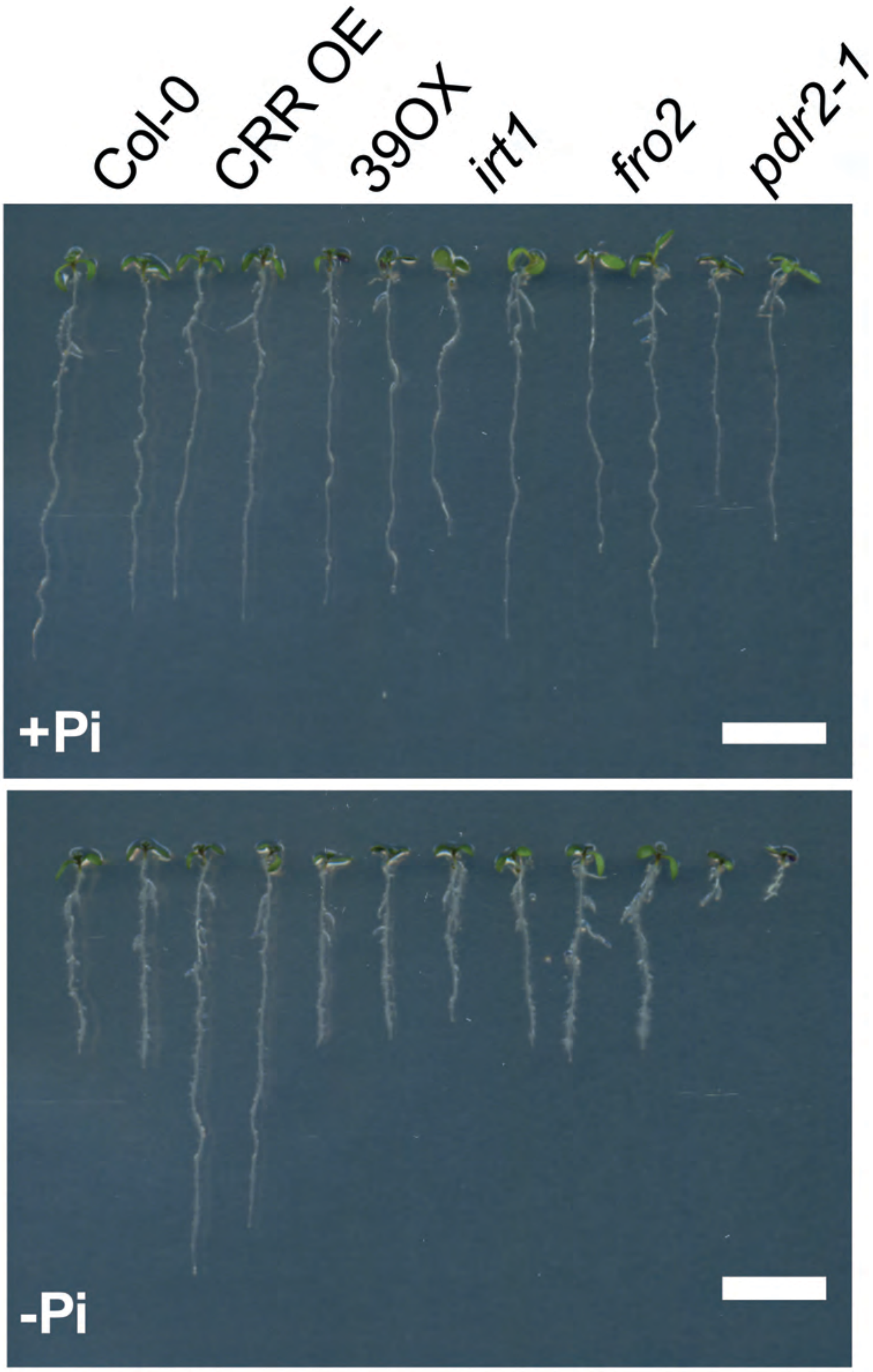


**Figure S11. Primary root growth of the iron homeostasis mutants *irt1* and *fro2* under Pi deficiency are similar Col-0.**
Photographs showing Col-0, CRR OE, 39OX, *irt*1, *fro2* and *pdr2* seedlings grown for 7 days under plus (+Pi) or minus (-P) phosphate conditions. Line 39OX over-express the transcription factor bHlH39 and shows a strong increase in root ferric-reductase activity. Scale: 1 cm.

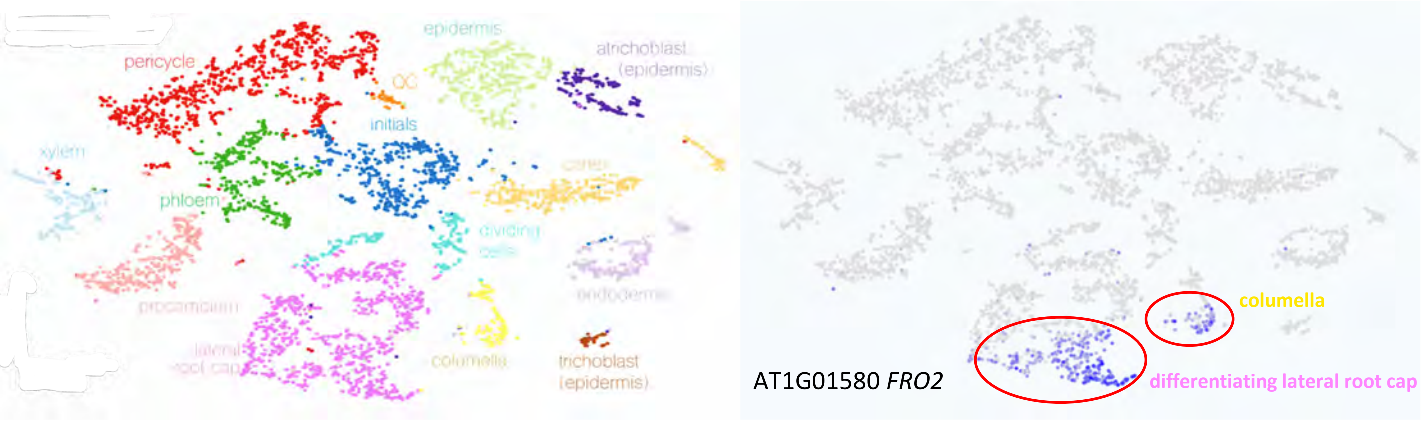


**Figure S12. *FRO2* expression pattern in root tips.**
Single-cell transcriptomic data from Arabidopsis root tips was retrieved from the Plant sc-Atlas (https://bioit3.irc.ugent.be/plant-sc-atlas/). The left panel shows a color-coded tSNE plot with the classification of sequenced cells into the different cell identities, and the right panel shows which cells express *FRO2*.

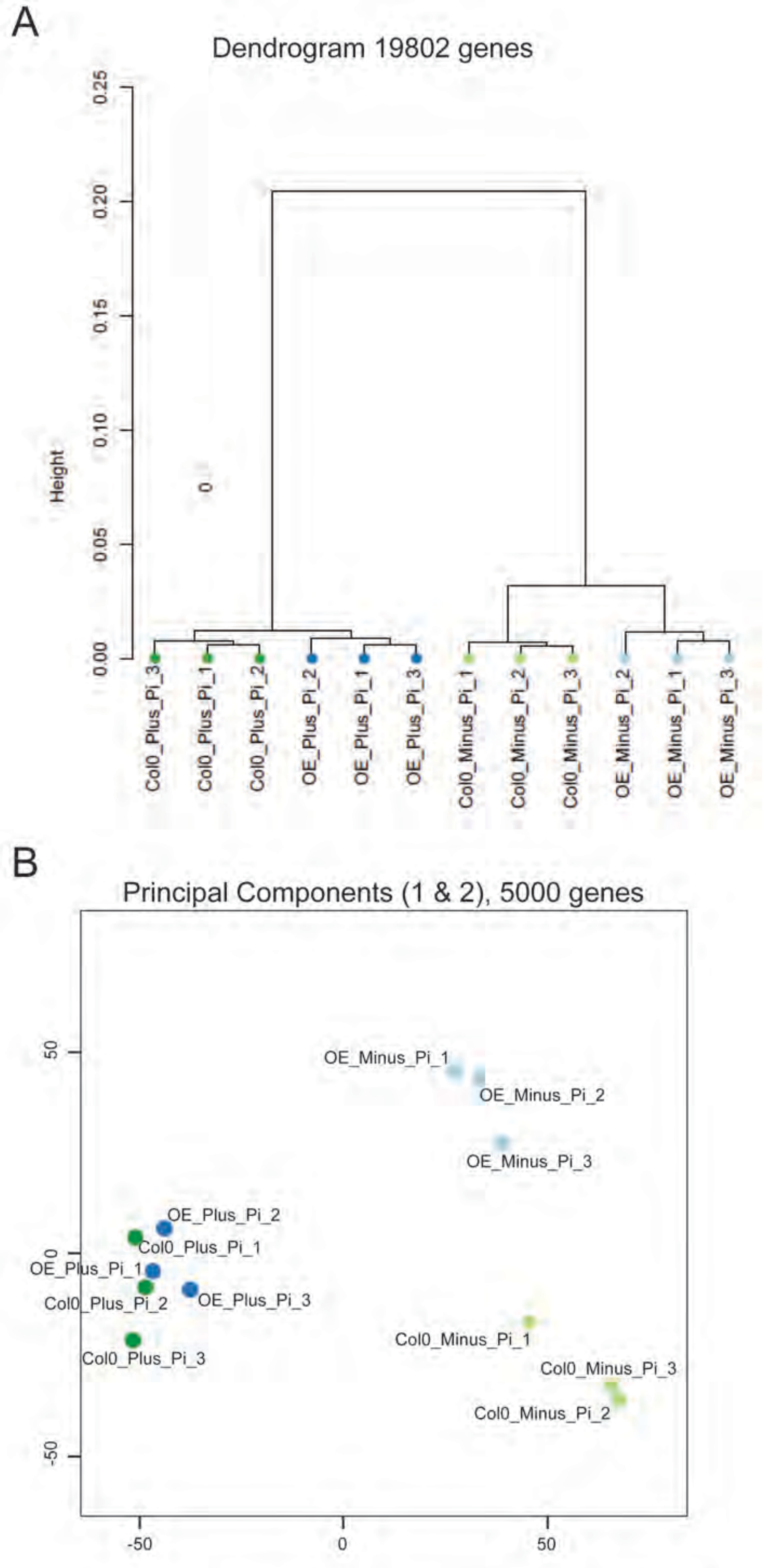


**Figure S13. Quality control and principal component analysis of the different RNA-seq samples used in the transcriptomic analysis**
**A,** Dendrogram showing that in all conditions the 3 biological replicates used in the RNA-seq analysis cluster together. **B**, Principal component analysis (PCA) based on the expression of the most 5000 variable genes. The PCA shows that Col-0 and CRR OE transcriptomes cluster together in plus phosphate (+Pi) conditions, while they diverge under minus phosphate (-Pi) conditions.

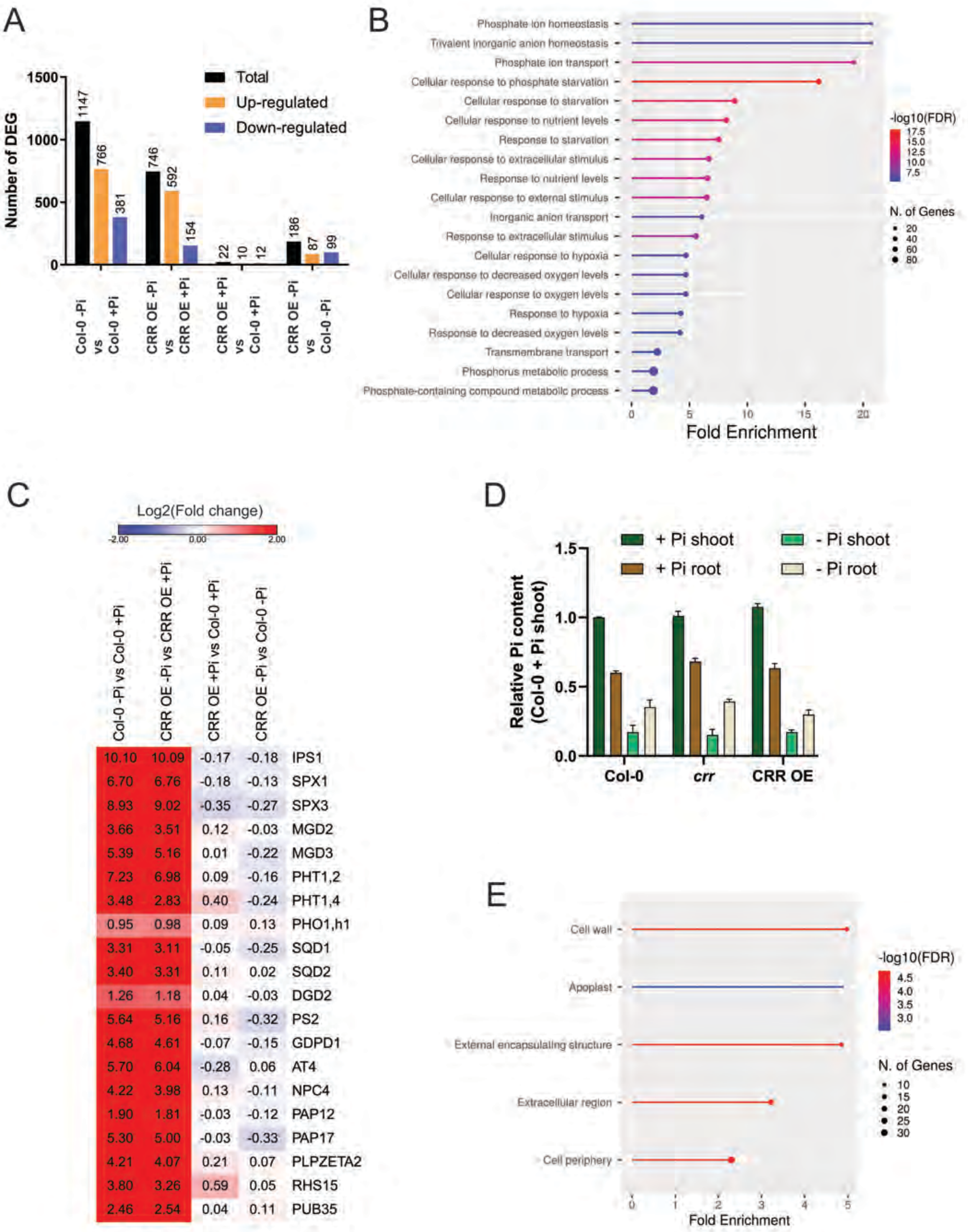


**Figure S14. Analysis of the differential expressed genes between Col-0 and CRR OE roots grown in +Pi and -Pi conditions**

**A**, Graph showing the total, up- and down-regulated number of differential expressed genes (DEG) found in the pairwise-comparisons indicated in the x-axis. **B**, Gene ontology (GO) enrichment analysis focused on molecular functions for DEG found the -Pi/+Pi treatments and shared by both Col-0 and CRR OE. The lollipop graph shows for each category the fold enrichment, the number of genes (N. of genes) and the -log10 of the fold discovery rate (FDR). **C**, Heat map showing the expression pattern of canonical phosphate starvation responsive genes. Data is expressed as the log2 of the fold change. **D**, Inorganic phosphate quantification of Col-0, *crr* and CRR OE roots and shoots from seedlings grown in plus (+Pi) or minus (-Pi) phosphate conditions. Data represented as mean ± s.d. After One-way ANOVAs analysis no significant differences were found between the genotypes analyzed. **E**, GO enrichment analysis focused on cellular compartment categories. The DEG found in the interaction between the genotype and treatment was used as input. The lollipop graph shows for each category the fold enrichment, the number of genes (N. of genes) and the -log10 of the FDR.

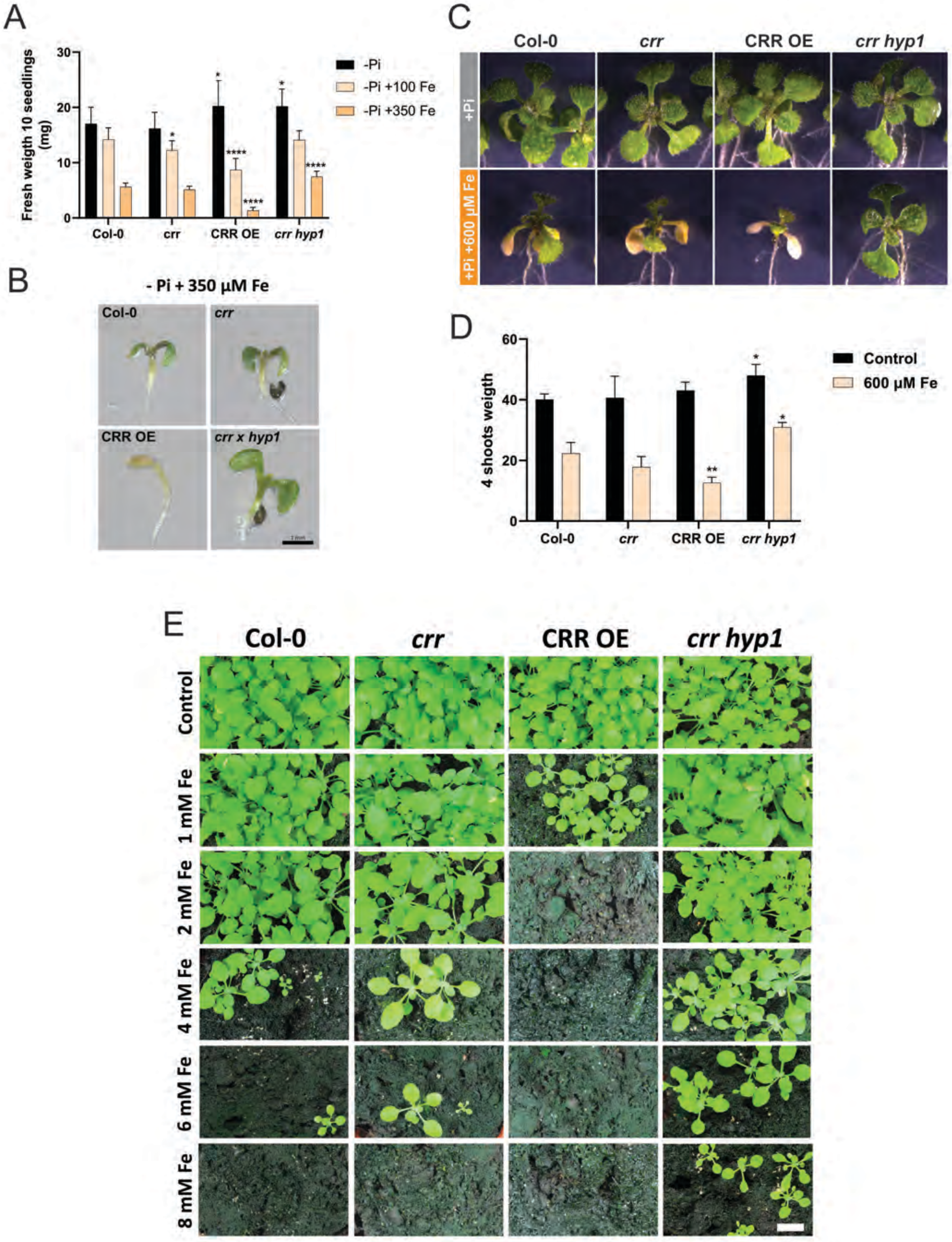


**Figure S15. The CYBDOM proteins CRR and HYP1 are involved in iron toxicity tolerance.**
**A**, Effect of iron toxicity under phosphate starvation conditions. Col-0, crr, CRR OE and *crr hyp1* double mutant were grown in minus phosphate (-Pi) conditions with the addition of increasing amounts of Fe-EDTA. Seven days after germination the fresh weight of 10 seedlings was recorded. **B**, Representative images showing the shoot phenotypes of seedlings grown in -Pi and 350 µM of Fe-EDTA. **C**, Effect of high iron levels on 7 days old seedlings. Plants were grown for 7 days in +Pi and then transferred to plates containing either the same media as a control or additional iron (600 µM Fe-EDTA). Four days after being transferred the shoots were photographed. **D**, Quantification of shoot biomass from the plants described in (C), expressed as the weight of 4 shoots (mg). In (A) and (D) data is represented as mean ± s.d (n≥3) and the statistical differences were analyzed using a One-way ANOVA followed by a Tukey's multiple comparisons test. Significant differences against Col-0 are indicated (***P < 0.001; **P < 0.01; *P < 0.05). **E**, Effect of iron toxicity on plants grown in soil. Seeds were sown in soil watered with tap water supplemented with increasing concentrations of Fe-EDTA. After 15 days the pots were photographed. Scale: 1 cm.

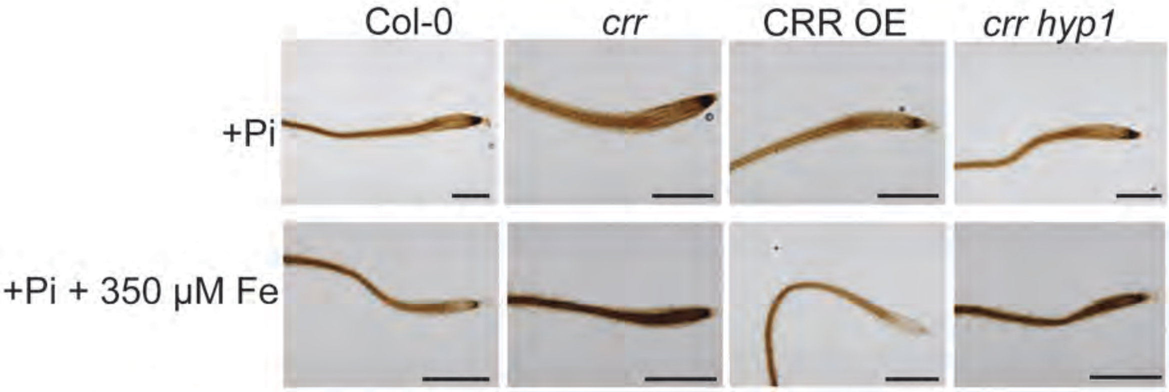


**Figure S16. CRR levels modulates the amount of iron accumulated at the root tip under iron stress.**

Representative images showing iron distribution pattern at the root tip of Col-0, *crr*, CRR OE and *crr hyp1* seedlings grown under plus phosphate (+Pi) conditions,  and +Pi with a supplement of 350 uM Fe-EDTA. Scale : 500 µm
